# RNA structural complexity dictates its ion atmosphere

**DOI:** 10.1101/2025.05.19.654961

**Authors:** Hiranmay Maity, Peter Zhang, D. Thirumalai, Hung T. Nguyen

**Affiliations:** Department of Chemistry, The State University of New York at Buffalo, NY, USA; Department of Chemistry, The University of Texas at Austin, TX, USA

## Abstract

Electrostatic interactions mediated by the surrounding ions govern virtually every facet of RNA behavior. Most studies have focused on rigid, well-folded motifs, leaving the structurally heteregeneous, and biologically ubiquitous, flexible RNAs underexplored. To address this gap, we performed molecular dynamics simulation of three RNAs spanning the structural continuum: an unstructured poly-uridylic tract (rU_30_), a semiflexible cytosine-adenine-guanine (CAG) repeat, and the tightly folded Beet Western Yellow Virus (BWYV) pseudoknot. Despite carrying nearly identical net charge, their ion atmospheres diverge strikingly. rU_30_ envelops itself in a diffuse Mg^2+^ environment retained through two or more hydration shells, whereas the CAG repeat and pseudoknot favor more localized outer-sphere Mg^2+^ binding. In contrast, Ca^2+^ tends to form inner-sphere contacts with all three RNAs, regardless of their folds. Remarkably, the diffuse ion clouds around unstructured RNAs extend farther into solution than that of the folded RNAs, significantly broadening their electrostatic sphere of influence. Nonetheless, the ion exchange kinetics remain virtually unchanged, demonstrating a surprising decoupling between spatial distribution and dynamical turnover. Our findings reveal RNA structural flexibility as a powerful lever for tuning ionic screening, with important implications for biomolecular recognition, RNA-driven phase separation, and physical properties of RNA-rich condensate.

**TOC Graphic:** 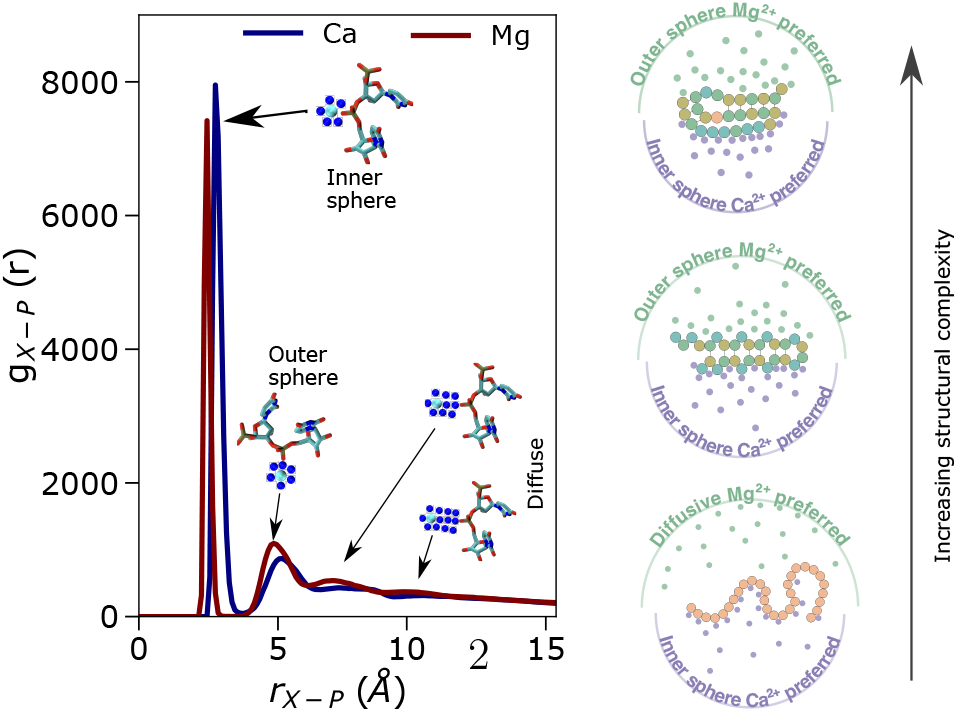

## Introduction

Metal ions are essential in assembling functional RNA structures ^1–4^ by neutralizing the negatively charged phosphate backbone, driving compaction, and controlling conformational changes in catalytic sites. ^5–7^ They also enable the formation of RNA condensates, ^8–11^ which can arise solely from homotypic interactions of low-complexity RNA sequences, such as polyuridylic acids (polyU) and cytosine-adenine-guanine (CAG) repeat sequences, in the absence of other biomolecules such as proteins. While both monovalent and divalent ions serve as counterions, RNA structure and condensate formation are highly sensitive to ion concentration, valency and identity. ^9,11–13^ Consequently, elucidating RNA-ion interactions remains central to experimental and theoretical studies on RNA folding ^14–29^ and phase separation.^8–12,30–39^

According to the Oosawa–Manning counterion condensation theory and non-linear Poisson– Boltzmann (PB) framework, ^40,41^ divalent cations stabilize RNA tertiary folds more efficiently than monovalent ions due to the reduced number of ions needed for charge compensation.^42,43^ Among divalent ions, Mg^2+^ is especially critical because its small radius and high charge density favor RNA tertiary structure formation, as seen in *Tetrahymena* ribozyme catalysis, which requires Mg^2+^, or to some extent Mn^2+^, but not others.^7,44^ Both simulations and experiments have demonstrated that ions bind nucleotides in a site-specific manner rather than uniformly,^45–54^ with divalent ions either partially dehydrated (inner-sphere, IS) or fully hydrated (outer-sphere, OS).^18^ Ions located beyond two hydration shells of the phosphate groups are classified as diffuse ions. A schematic illustrating IS, OS, and diffuse ions surrounding an RNA segment is shown in Fig. 1A.

**Figure 1:**
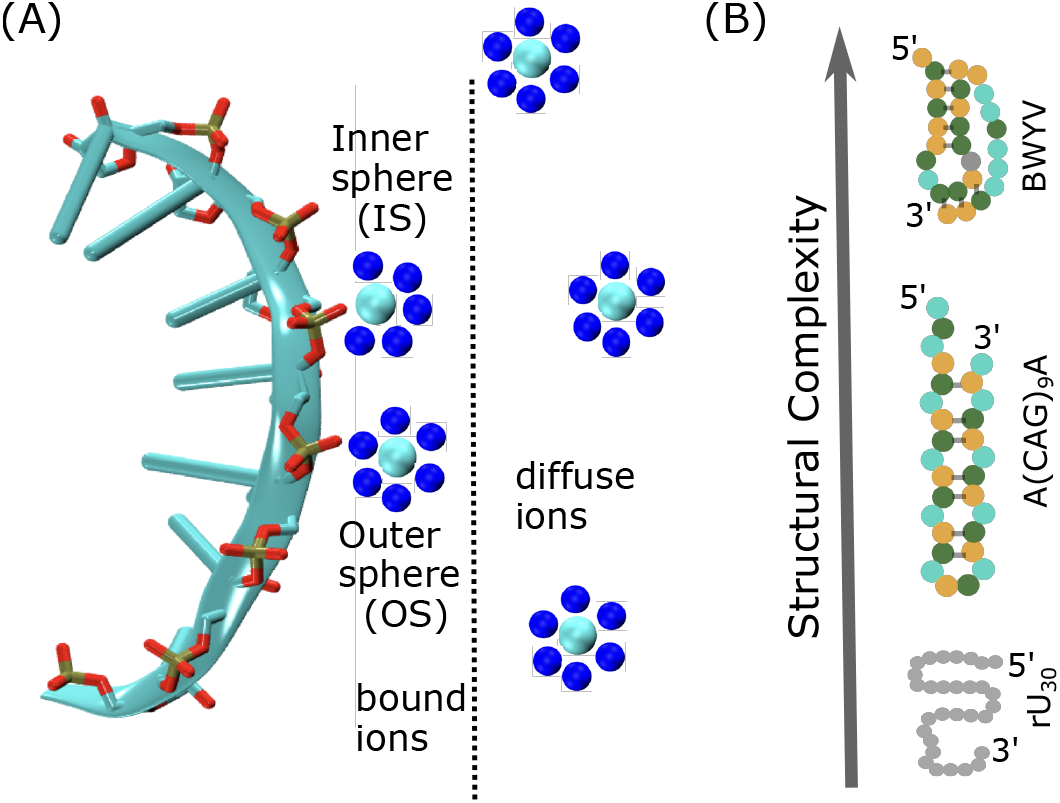
Illustration of binding modes of Mg^2+^ and RNA structural complexity: (A) Mg^2+^ ions can bind directly to RNA without intervening water molecules (inner-sphere, IS), or interact via one or multiple water molecules, referred to as outer-sphere (OS) and diffuse ions, respectively. (B) Schematics of three different types of RNA with distinct structural features. Homopolymeric sequences such as rU_30_ lack base pairs and adopt random coil-like conformations. Repeat RNAs such as CAG repeats exhibit stem-loop or hairpin-like conformations. BWYV RNA forms a pseudoknot with stable secondary and tertiary structures.

Extensive research has investigated the ion atmosphere surrounding well-structured RNAs that adopt unique three-dimensional conformations under ambient conditions. However, flexible RNA sequences that exhibit significant structural heterogeneity and are prone to condensate formation remain less explored. Recent experiments on single-stranded nucleic acids reveal notable differences in their ion atmospheres, but offer only a coarse view of RNA-bound ions.^55–57^ How the ion atmosphere changes with varying degrees of RNA structural flexibility remains underexplored.

In this study, we examine three RNAs of similar net charge but differing structural complexity (Fig. 1B): (i) 30-nucleotide polyuridylic acid (rU_30_), which is highly flexible;^58^ (ii) a 9-repeat cytosine-adenine-guanine (CAG) RNA, A(CAG)_9_A, which adopts multiple stem-loop conformations; ^59^ and (iii) the 28-nucleotide beet western yellow virus (BWYV) pseudoknot, which possesses a stable tertiary fold.^60^ We simulate those RNAs using the Single Interaction Site (SIS) model, which has been successfully used to study the structures and phase behaviors of repeat RNAs^30,59,61,62^ as well as interactions and oligomerization of mRNAs.^63–65^ In our simulations, RNAs are modeled in a mixture of monovalent and divalent ions, X^2+^ (X is either Mg or Ca), using a recently developed method^66,67^ based on the Reference Interaction Site Model theory^68,69^ to accurately capture the RNA structural ensembles and ion distributions.

Our results reveal that ion binding modes and their populations depend strongly on RNA structural complexity, ion concentration, and cation type. Unstructured rU_30_ favors diffuse Mg^2+^, whereas structured RNAs exhibit pronounced Mg^2+^ OS binding. We showed that ion distributions around rU_30_ are uniform, while specific binding emerges in A(CAG)_9_A and BWYV, especially at low ion concentrations. Moreover, the cation charge density plays a key role in dictating the ion atmosphere as Mg^2+^ typically engages in OS contacts, while Ca^2+^ strongly prefers IS binding. Most importantly, we demonstrated that the ion atmosphere of unstructured rU_30_ extends further into solution compared to folded RNAs, despite having a fewer number of bound ions. Surprisingly, variations in RNA structure have negligible effects on ion dynamics, perhaps due to the simple nature of those RNAs. Overall, our results reveal that ion-RNA interactions are governed by a delicate interplay between ion properties and RNA architecture, which is itself stabilized by those same ions. Given the critical role of the ion atmosphere in protein binding thermodynamics and RNA phase separation propensity, these insights could guide the design of new biomaterials with tailored properties and phase behaviors.

## Results

### Ion effect on RNA conformational ensemble

To explore the effects of ions with different valency on RNA structural ensemble, we simulated RNAs in the mixture of monovalent salts and divalent cations. In our model, monovalent salts are treated implicitly, while divalent cations are represented explicitly (see Methods). This combination enables efficient sampling of the RNA conformational space while still preserving the RNA electrostatics and folding thermodynamics.^66,67,70^

#### rU_30_

We observed that rU_30_ undergoes a significant collapse as Mg^2+^ concentration is increased from 0 to 3 mM (Fig. 2A), in qualitative agreement with small angle X-ray scattering (SAXS) data.^55^ Interestingly, the radius of gyration, ⟨*R*_*g*_⟩, obtained at 20 mM NaCl + 1 mM Mg^2+^ is identical to one at 150 mM NaCl without Mg^2+^ (Fig. 2A, Fig. S1A), which is ≈ 29.1 Å. This suggests that the effect of Mg^2+^ could be equivalent to 100-fold of monovalent ions.^71^ Similarly, the average pair distance ⟨*R*_*ij*_⟩, as a function of the sequential separation length |*i* − *j*|, decreases substantially with an increase in Mg^2+^ concentration (Fig. S2B) (mostly by long-ranged interactions |*i* − *j*| ≥ 8). The computed contact probability with respect to a Gaussian polymer chain, ⟨*Q*_*ij*_⟩ (Eq. 3), for a given sequential separation length |*i* − *j*| increases with an increase in either Mg^2+^ (Fig. S2) or monovalent ion concentrations (Fig. S1C S1F). Nonetheless, the contact maps are featureless at all ion concentrations, signifying that the compaction is non-specific and the RNA conformational ensemble remains heterogeneous. Our results that rU_30_ samples a heterogeneous ensemble of conformations (Fig. 2A) are in accordance with recent experiments. ^55,58^ Similar conclusion also holds true in the presence of monovalent salt alone (detailed in Supporting Information (SI)).

**Figure 2:**
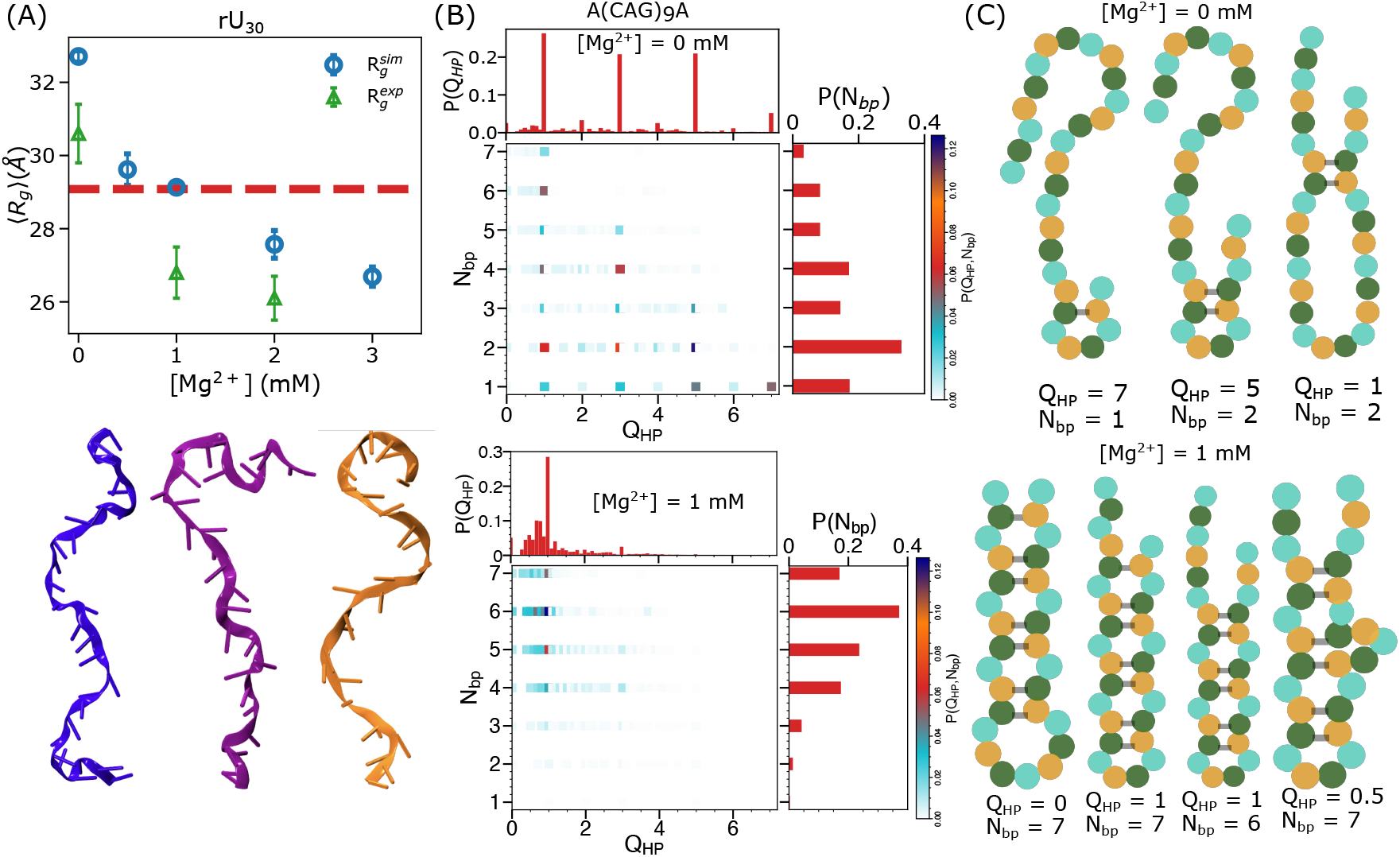
Structures of rU_30_ and A(CAG)_9_A: (A) (top) Comparison between simulated (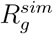, blue circles) and experimentally measured (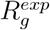, green triangles) radius of gyration for rU_30_ as a function of Mg^2+^ concentration at 20 mM NaCl. The red dashed line indicates 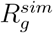 at 150 mM NaCl without Mg^2+^. (Bottom) Representative structures of rU_30_ at 20 mM NaCl + 1 mM Mg^2+^. All atom structures are generated from coarse-grained coordinates using Arena web server. ^74^ (B) Joint probability distributions of *Q*_*HP*_ and *N*_*bp*_, *P* (*Q*_*HP*_, *N*_*bp*_), for A(CAG)_9_A at 20 mM NaCl without (top) or with 1 mM Mg^2+^ (bottom) demonstrate that the CAG repeat adopts multiple stem-loop-like conformations. Marginal distributions of *Q*_*HP*_ (*P* (*Q*_*HP*_)) and *N*_*bp*_ (*P* (*N*_*bp*_)) are shown in the top and right hand sides, respectively. (C) Schematics of A(CAG)_9_A secondary structures at 20 mM NaCl without (top) or with 1 mM Mg^2+^ (bottom). Associated *Q*_*HP*_ and *N*_*bp*_ values are denoted below each structure.

#### CAG repeat

In contrast to rU_30_, the CAG repeat adopts multiple stem-loop-like conformations due to complementary base pair interactions. ⟨*Q*_*ij*_⟩ values for A(CAG)_9_A in both monovalent (Fig. S3) and Mg^2+^ ions (Fig. S4) exhibit the signature of stem-loop-like structures in the conformational ensemble. To quantify the RNA ensemble, we used the order parameter *Q*_*HP*_ (Eq. 2), which quantifies the arrangements of base pairs in the stem region.^59^ We constructed *Q*_*HP*_ in such a way that if *Q*_*HP*_ = *m* (*m* is a non-negative integer) then there is an overhang with *m* unit(s) of unpaired CAG repeat(s) at the terminal of the stem-loop or hairpin structure. On the other hand, a fractional value of *Q*_*HP*_ denotes a bulge in the stem region. Fig. 2B shows the joint probability distribution of *Q*_*HP*_ and *N*_*bp*_, where *N*_*bp*_ is the number of base pairs in the stem region. At 20 mM NaCl, the ensemble is heterogeneous and the RNA mainly populates a partially folded structure that has *N*_*bp*_ = 2 base pairs and *Q*_*HP*_ = 5 CAG repeats overhang at the terminal. Importantly, *Q*_*HP*_ exhibits significant populations of hairpins with more odd numbers of overhang (*m* = 1, 3, …) than even-numbered overhangs (*m* = 2, 4, …), which is a characteristic of odd-numbered CAG repeats.^59^ The RNA is mostly unfolded, which is reflected in the low number of *N*_*bp*_ (upper panel of Fig. 2C). Upon addition of 1 mM Mg^2+^, it folds to a stable hairpin that contains one CAG unit overhang (bottom panel of Fig. 2B, S6) where *Q*_*HP*_ = 1 and *N*_*bp*_ = 6, which is expected for the odd-numbered CAG repeats. ^59,72,73^ Notably, the CAG ensemble is significantly more collapsed compared to those in monovalent salt, because RNA now samples mostly folded hairpin structures for which the stem region contains at least 4 base pairs (bottom panel of Fig. 2C).

### Increased RNA structural complexity leads to enhanced ion accumulation

Due to the interplay between RNA conformational ensemble and ion-RNA interactions, we expected that the ion atmosphere surrounding RNA would depend on its structural complexity. In addition to rU_30_ and the CAG repeat for which the RNA adopts multiple structures at ambient conditions, we also considered the BWYV pseudoknot which adopts a stable secondary and tertiary structure. We deliberately chose those RNAs with similar net charges to focus on the effect of RNA structural flexibility alone. We performed coarse-grained simulations of BWYV by restraining it around the crystal structure (PDB ID 437d), allowing only ions to equilibrate during simulations. The radial distribution function *g*_Mg*−*P_(*r*) between the phosphate groups and Mg^2+^ displays multiple peaks, each representing distinct solvation shells of ions around RNA, before converging to the bulk level at large separation (Fig. 3A).

**Figure 3:**
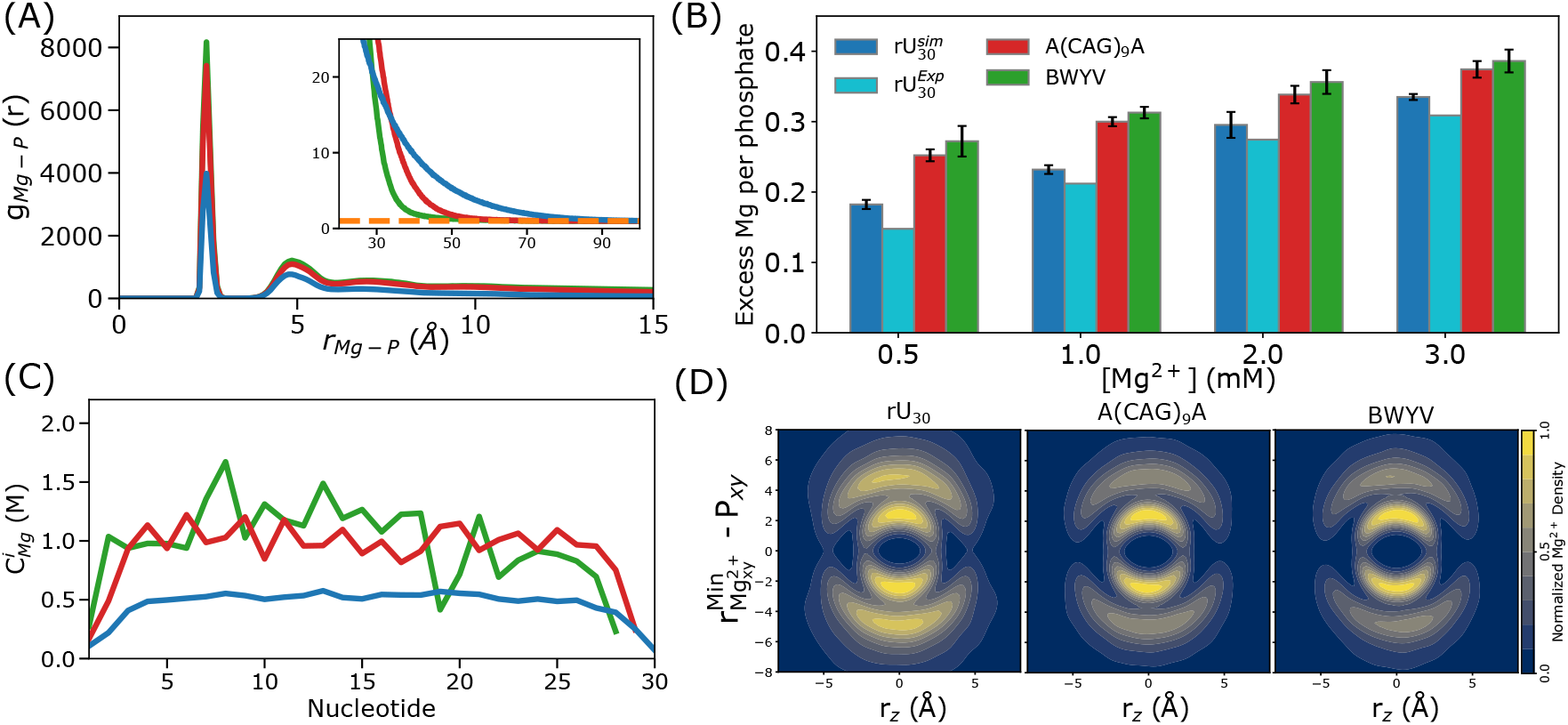
Ion distribution around RNAs: (A) Radial distribution function *g*_Mg*−*P_(*r*) between Mg^2+^ and phosphate groups for rU_30_, A(CAG)_9_A and BWYV. *g*_Mg*−*P_(*r*) decays rapidly for well-structured RNAs (inset). The orange dotted line (inset) denotes the bulk level of Mg^2+^ for which *g*_Mg*−*P_(*r*) = 1. (B) Excess Mg^2+^ per phosphates for rU_30_, A(CAG)_9_A and BWYV. The experimental values^55^ for rU_30_ (cyan) are determined at 4°C. (C) Local Mg^2+^ concentrations for rU_30_, A(CAG)_9_A, and BWYV are correlated with the phosphate density. (D) 2D spatial distribution of Mg^2+^ with respect to the closest phosphate group for rU_30_ (left), A(CAG) A (middle) and BWYV (right). 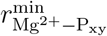 and *r*_*z*_ are the minimum distances between the phosphate and Mg^2+^ on the xy-plane, and along the z-plane, respectively (see Materials and Methods).

The magnitudes of the first and second peaks decrease with RNA structural complexity, following the order BWYV *>* A(CAG)_9_A *>* rU_30_ (Fig. 3A), indicating that the ions preferentially bind in the proximity of structured RNAs. This is also reflected in the excess number of ions condensed onto the phosphate groups Γ_*Mg*_ ^55,75,76^ (Fig. 3B), calculated using the Kirkwood–Buff integral (Eq. 4).^77^ Γ_*Mg*_ increases with elevating Mg^2+^ concentrations and more importantly, increases with RNA structural complexity. BWYV, which adopts a compact pseudoknot structure with a high phosphate density, shows a much higher excess number of Mg^2+^ ions per phosphate group.^60,66^ In contrast, rU_30_, an unstructured RNA, exhibits a lower excess number of Mg^2+^ despite carrying a similar net charge. ^55^ The CAG repeat, which forms multiple stem-loop-like conformations, shows an excess number of Mg^2+^ that is intermediate, sandwiched between BWYV and rU_30_. This is because the higher charge density of folded RNAs compared to unstructured counterparts favors enthalpic interactions with ions. Our finding is consistent with the expectation from Oosawa–Manning ion condensation theory, ^40^ classical PB electrostatics, numerous experimental measurements and computer simulations. ^66,75^

### Specificity of divalent cation binding

To investigate whether condensation of ions onto RNA occurs uniformly, we computed the local Mg^2+^ concentration, 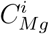 (Eq. 5), at individual nucleotide positions for three RNAs at 1 mM Mg^2+^ (Fig. 3C). Interestingly, 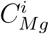 shows that ion condensation could be both site specific and non-specific depending on the RNA structure. rU_30_, lacking a stable secondary structure and thus sampling an ensemble of rapidly interconverting structures, binds Mg^2+^ in a non-specific manner. In contrast, BWYV pseudoknot exhibits a fingerprint for specific Mg^2+^ binding sites. The CAG repeat, on the other hand, samples multiple hairpin-like structures, therefore bearing a weak signature of site specific Mg^2+^ condensation due to its repeat sequence. Nucleotides in close proximity in the stem region of the CAG hairpin attracts divalent cations more strongly than the loop region, which is in agreement with previous simulations.^45,67^ Interestingly, the site-specific binding becomes less prominent at high ion concentrations (Fig. S7). Our observations remain valid when Ca^2+^ is used instead of Mg^2+^ (Fig. S8).

### Ion atmosphere expands in flexible RNAs

Due to the stronger ion interactions with folded RNAs compared to unstructured counterparts, we expected that the ion atmosphere of folded RNA should extend further in space. Surprisingly, we found the opposite result, *i*.*e*. the distance at which the ion concentration converges to the bulk value (*g*_Mg*−*P_(*r*) → 1) follows the reverse order: BWYV *<* A(CAG)_9_A *<* rU_30_ (inset Fig. 3A). To further illustrate the ion atmosphere, we projected the ion distribution onto a 2D density plot after aligning RNA along its principal axes (Fig. 3D). This analysis clearly reveals a collapse of the ion atmosphere as the RNA structural complexity increases. Such collapse is also apparent in the radial distribution function of Mg^2+^ to the RNA center of mass (Fig. S9), and the histogram of Mg^2+^ to the closest phosphate group (Fig. S10).

To understand the reason, we further partitioned the ion atmosphere into IS, OS and diffuse regions (illustrated in Fig. 1A), depending on the ion-P distance (IS: *r*_Mg*−*P_ ≤ 3.2 Å, OS: 3.2 Å*< r*_Mg*−*P_ ≤ 6.1 Å). The number of diffuse ions is then calculated as *N*_*D*_ = Γ_*X*_ −*N*_*IS*_ −*N*_*OS*_, where Γ_*X*_ is the excess number of ions, and *N*_*IS*_ and *N*_*OS*_ are the numbers of IS and OS ions, respectively. We observed that in general, there is enhancement of all binding modes at elevated Mg^2+^ concentrations, albeit with varying degrees (Fig. 4A). IS binding is less effective than OS binding (Fig. 4C, S11A) for those three RNAs, which is consistent with previous studies.^67,78^ This is likely due to the high entropic cost associated with localizing ions in the RNA proximity. Notably, OS ions contribute the most for folded RNAs (CAG repeat and BWYV), while the diffuse region is dominant for flexible and unstructured rU_30_ (Fig. 4A). Thus, the ion atmospheres of unstructured RNAs extend further, reaching beyond those of stable and well-folded RNAs. Distant ions, located far from the RNA yet still interacting with it due to long range electrostatics, serve as the RNA’s spatial fingerprint and thus can be crucial for its interactions with other biomolecules. Our finding that unstructured RNAs possess a more extensive ion atmosphere, and thus influence a broader region of the surrounding solution, may have implications for their roles in molecular assembly and biological function.

**Figure 4:**
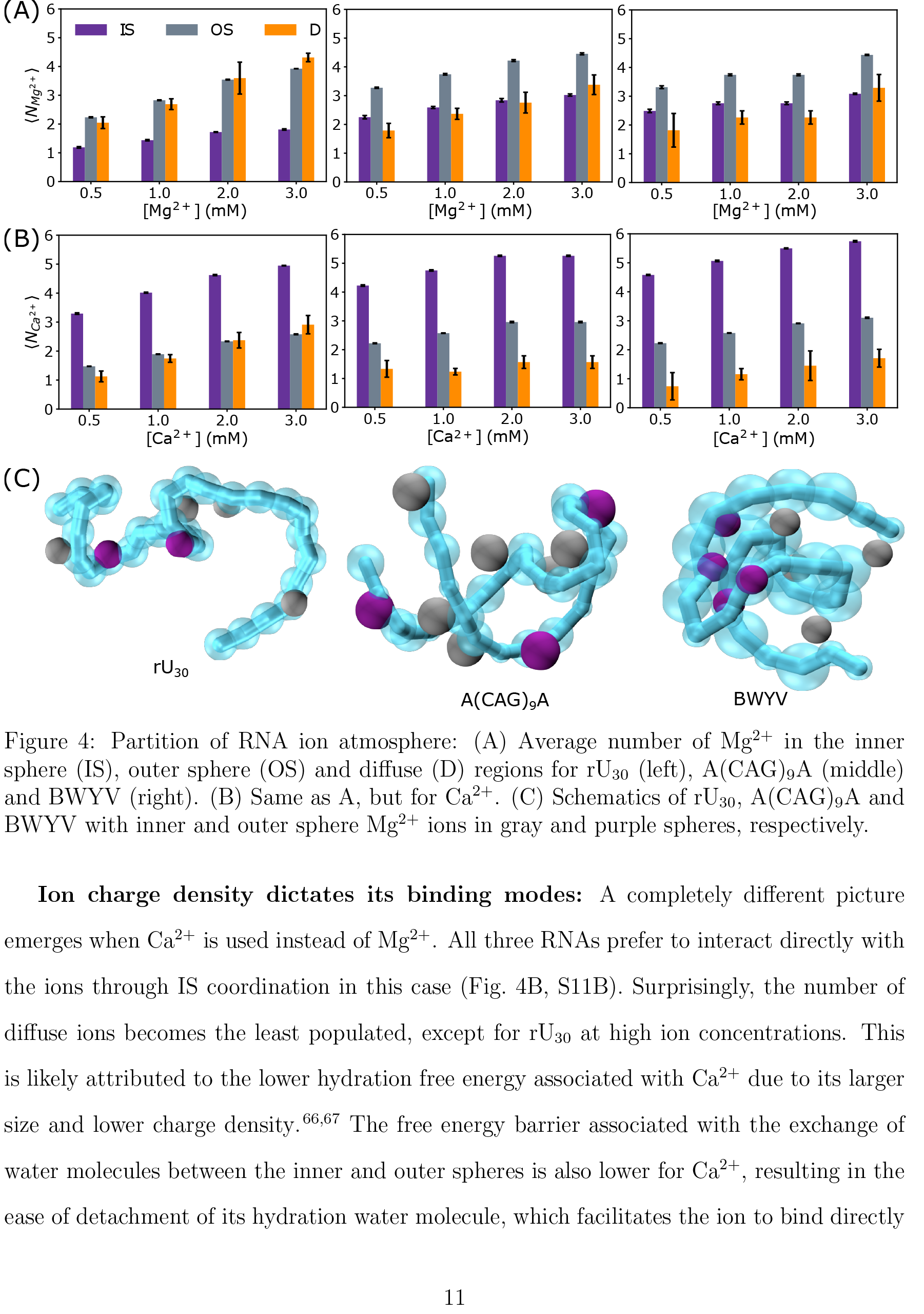
Partition of RNA ion atmosphere: (A) Average number of Mg^2+^ in the inner sphere (IS), outer sphere (OS) and diffuse (D) regions for rU_30_ (left), A(CAG)_9_A (middle) and BWYV (right). (B) Same as A, but for Ca^2+^. (C) Schematics of rU_30_, A(CAG)_9_A and BWYV with inner and outer sphere Mg^2+^ ions in gray and purple spheres, respectively.

### Ion charge density dictates its binding modes

A completely different picture emerges when Ca^2+^ is used instead of Mg^2+^. All three RNAs prefer to interact directly with the ions through IS coordination in this case (Fig. 4B, S11B). Surprisingly, the number of diffuse ions becomes the least populated, except for rU_30_ at high ion concentrations. This is likely attributed to the lower hydration free energy associated with Ca^2+^ due to its larger size and lower charge density.^66,67^ The free energy barrier associated with the exchange of water molecules between the inner and outer spheres is also lower for Ca^2+^, resulting in the ease of detachment of its hydration water molecule, which facilitates the ion to bind directly to phosphate groups.^79^ The shift in ion binding from long range with Mg^2+^ to short range with Ca^2+^ may influence Nature’s selection of specific ions based on biological context and functional needs.

### Ion exchange dynamics is insensitive to RNA folds

We now turn to the question of how dynamics of the ion atmosphere is affected by the RNA conformational flexibility. To characterize ion dynamics, we conducted Brownian simulations to calculate the dwell time of bound ions in the IS 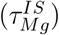 and OS 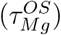 regions, shown in Figs. 5A and 5B, respectively, at 20 mM NaCl + 1 mM Mg^2+^. These distributions exhibit exponential decay, indicating a single characteristic timescale associated with the ion’s dwell times. We extracted such timescales, *τ*_0_, by fitting the dwell time distribution to *c*_0_ exp(−*τ/τ*_0_). Despite clear differences in the ion atmosphere for different RNA constructs, we did not observe any significant changes of the ion relaxation for those RNAs. For IS ions, *τ*_0_ in rU_30_, CAG repeat, and BWYV are 4.1 ns, 3.6 ns, and 4.1 ns, respectively (Fig. S12A). For OS ions, the relaxation time is more than an order faster, with *τ*_0_ values being 92 ps, 125 ps, and 115 ps, respectively (Fig. S12B). As expected, the timescale of IS residence is significantly longer than the OS counterpart due to the much stronger IS binding. Our dwell time estimates are in close agreement with experimental measurements and all-atom simulations. ^80,81^ We also found a negligible dependence of ion dwell times on Mg^2+^ concentrations (Figs. 5C and 5D). Nevertheless, the ion atmosphere remains highly dynamic (Fig. S13), with ions exhibiting nearly uniform average dwell times around individual nucleotides (Fig. S14).

**Figure 5:**
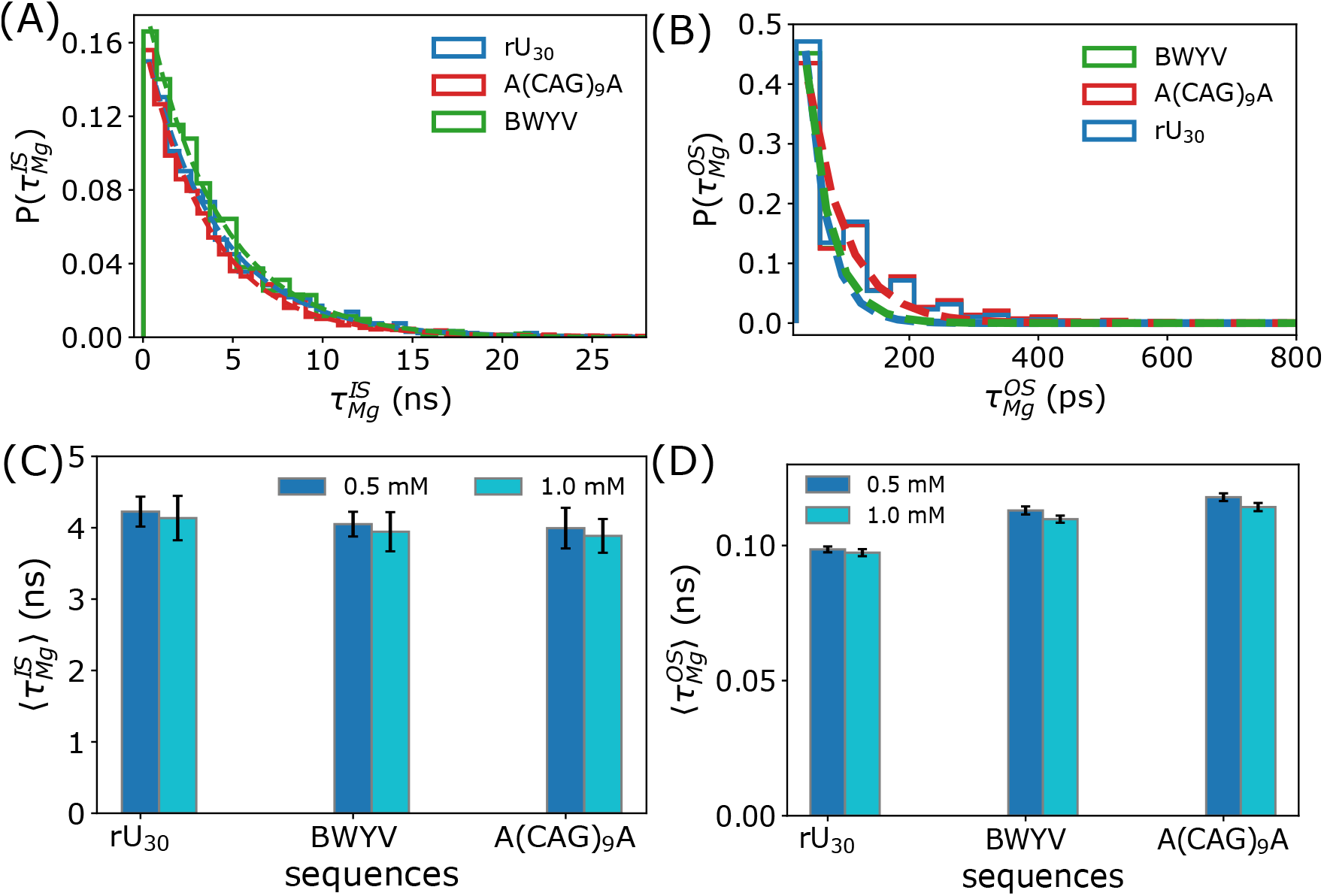
Dwell time of bound Mg^2+^ ions: (A) Probability distribution of the dwell time of ion in inner sphere (IS) region for rU_30_, A(CAG)_9_A and BWYV. Dashed lines indicate single exponential fits to the dwell time distribution. (B) Same as A, but for outer sphere (OS) region. (C) Average dwell time of IS ions. (D) Average dwell time of OS ions.

Our observation that the dwell time is independent on the RNA fold is likely due to the simple nature of RNAs studied here, which are relatively small and thus do not fold into complex architecture with deep binding pockets. In other more complex RNAs with such binding sites and high electrostatic potentials, ions could easily get stuck for a long time and become an integral part of the RNA structure, which are frequently detected in X-ray crystallography structures.

## Conclusion

Electrostatic interactions play a dominant role in stabilizing RNA structures and dictate their interaction with neighboring molecules. In this work, we investigated the ion atmosphere in the RNA vicinity using coarse-grained molecular dynamics simulations. Our results reveal that ion binding modes and their relative contributions depend strongly on both ion concentrations and RNA structural flexibility. For unstructured RNAs such as rU_30_, diffuse ion populations are predominant. In contrast, for RNAs with defined secondary and tertiary structures, such as CAG repeats or BWYV pseudoknot, direct coordination, either by IS or OS, becomes dominant. Consequently, structurally flexible RNAs extend their electrostatic influence farther into solution, which may enhance their ability to engage with nearby molecular partners.

We also demonstrated that the counterion identity plays a key role in shaping the RNA ion atmosphere. When Ca^2+^ is used instead of Mg^2+^, IS binding becomes important, leading to a significant collapse of the ion atmosphere. Because of that, Ca^2+^ can compensate the negative charges of the phosphate groups more effectively than Mg^2+^, presumably leading to a higher propensity to induce RNA condensation. ^11,82^ Nonetheless, we observed that ions bind transiently to the RNA backbone and exhibit a high degree of mobility. During the complexation process involving two oppositely charged intrinsically disordered proteins, these bound ions are readily released into the bulk solution, ^83,84^ thereby favorably contributing to the entropy gain of the complexation process. Due to their transient dwell time near the RNA, divalent cations can similarly be released into the bulk during the complexation of RNA with oppositely charged proteins.

These findings have broad implications because the composition and extent of the ion atmosphere, dictated by RNA structural complexity, directly impact the electrostatic and physicochemical properties of RNAs in both dilute and condensed phases.^85^ For example, unstructured RNAs, enriched in diffuse and weakly interacting ions, have been shown to readily form liquid-like condensates. ^10,86^ The dynamic and weakly bound nature of these ions also enables them to mediate interactions with other biomolecules by being readily displaced during complexation. RNAs that sustain an extended diffuse ion cloud may act as long range recruiters of protein partners. In contrast, compact ionic shells associated with highly folded RNAs or with Ca^2+^ may promote local compaction and structural transitions that bias toward aggregation or scaffold organization.

## Materials and Methods

### Models

Following our earlier studies, ^30,59^ we represent each nucleotide by a single bead. In this work, we further establish that the effects of divalent cations could be taken into account within our framework. In the simulations, the monovalent salts are mimicked by the Debye–Hückel (DH) approximation, whereas the divalent cations are present explicitly. ^66^ The DH interactions are modeled using:

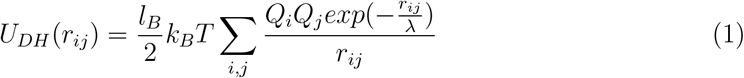

where *λ* = (8*πl*_*B*_*ρ*_1_)^*−*1*/*2^ is the Debye length that depends on the Bjerrum length 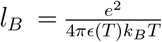. *e* is the elementary charge, *ϵ*(*T*) is the temperature dependent dielectric constant of water, *k*_*B*_*T* is the thermal energy and *ρ* is the number density of the monovalent salt. For divalent cations *Q*_*i*_ = 2, while the phosphate charge is renormalised to *Q*_*P*_ (*C*_1_, *C*_2_, *T*) that depends on the concentration of monovalent salt *C*_1_, divalent ions *C*_2_, and temperature *T*. The expression of *Q*_*P*_ (*C*_1_, *C*_2_, *T*), accounting for the condensation of monovalent salt onto the phosphate group, is given elsewhere.^66^ The T-dependence of the dielectric constant^87^ is *ϵ*(*T*) = 87.74 - 0.4008*T* + 9.398 × 10^*−*4^ *T* ^2^ - 1.410 × 10^*−*6^ *T* ^3^ (T expressed in °C). The detailed description of the energy functions of the RNA model is given in the SI.

### Criterion for base pair formation

A base pair between two complementary nucleotides, *I* and *j* is formed if their hydrogen bond energy 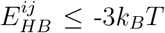, where *k*_*B*_ is the Boltzmann constant and *T* is the absolute temperature.

### Structural order parameters, Q_HP_

To characterize the hairpin structures adopted by the CAG repeat, we use the order parameter expressed as,^59^

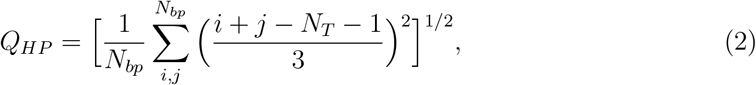

where *i* and *j* are the nucleotides base-pairing to each other, *N*_*T*_ is the sequence length, and *N*_*bp*_ is the number of base pairs in a given conformation. For a perfect hairpin, it is easy to show that *i* + *j* = *N*_*T*_ + 1, implying that *Q*_*HP*_ = 0. The value of *Q*_*HP*_ = 1 corresponds to a slipped hairpin with one unit of unpaired CAG. Fractional values, 0 *< Q*_*HP*_ *<* 1, represent a variety of conformations containing bulges in the stem.

### Contact probability, Q_ij_

We calculated the probability of contact formation between a pair of nucleotides *i* and *j* using,

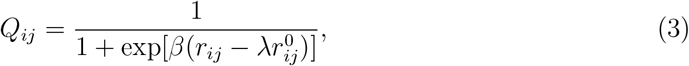

where *r*_*ij*_ represents the distance between the beads *i* and *j*, while 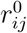 is the cutoff distance at which the probability of contact formation is 0.5. Using the Gaussian polymer chain as a reference system, the value of 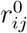 is taken as 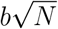, where *b* = 5.9 Å is the average bond length between two phosphates, and *N* = |*i* − *j*|− 1 is the number of bonds connecting beads *i* and *j*. We use *λ* = 1.5 to account for thermal fluctuation. We set *β* = 2.0 Å^*−*1^, which controls the rate at which the probability of contact decreases from 1 to 0.

#### Excess number of Mg^2+^, Γ_Mg_

To compute Γ_*Mg*_ around the RNA, we use the Kirkwood-Buff integral,^77^

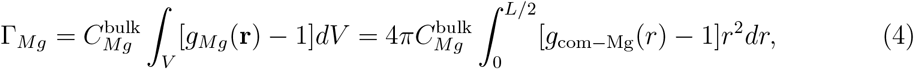

where 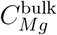 is the bulk concentration of the ions, and *g*_com*−*Mg_(*r*) is the radial distribution function between the center-of-mass (COM) of the RNA and Mg^2+^ ions. To ensure convergence, we integrate up to half of the simulation box length, *L/*2. *L* ≈ 720 Å for 1 mM ≤ [Mg^2+^] ≤ 3 mM and *L* ≈ 900 Å at 0.5 mM Mg^2+^. The excess number of Mg^2+^ per phosphate is computed as Γ_*Mg*_*/N*_*T*_, where *N*_*T*_ is the number of phosphate groups in the RNA.

*Local X*^*2+*^ *(X = either Mg or Ca) concentration around individual phosphate groups*, 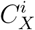:

We calculated 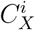 as,

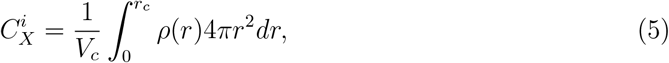

where *V*_*c*_ is the volume of a sphere with radius *r*_*c*_, *ρ*(*r*) is the density of ions at a distance *r* from a specific phosphate group. The cutoff distance, *r*_*c*_ = 3.2 Å for Mg^2+^ and 3.7 Å for Ca^2+^, accounts for the IS interaction of ions with the phosphate group.

#### 2D spatial distribution of Mg^2+^

We first prepare each snapshot by removing the translational and rotational movements of the RNA, where the RNA COM and principle axes are aligned. Subsequently, for each phosphate group, we calculated the distance to the nearest Mg^2+^ ion along the z-axis (*r*_z_), and its distance in the xy-plane 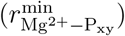, representing the shortest lateral separation. To further characterize the angular distribution, we computed the azimuthal angle, *ϕ* = arctan ^*y*^, where *x* and *y* are the in-plane components of the vector from the phosphate to the Mg^2+^ ion. The sign of the lateral distance in the xy-plane was then determined using sgn(*ϕ*), leading to 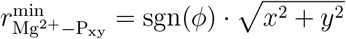.

#### Dwell time, τ

We computed *τ* of ions associated with the IS 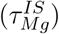 and OS 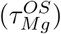 by calculating the time difference between ion entry and exit: *τ* = *t*_*out*_ − *t*_*in*_.

#### Simulations

Details of the simulations are identical to our previous work, and are reported in the SI.^59^ All thermodynamic properties are calculated using trajectories generated by integrating the Langevin equation of motion in the low friction limit using OpenMM. ^88^ To capture the ion dynamics, we used friction coefficient equivalent to water and applied BrownianIntegrator available in OpenMM.

## Acknowledgements

We gratefully acknowledge support for this work from the University at Buffalo to HTN. Computational support is provided by the Center for Computational Research at the University at Buffalo and Pittsburgh Supercomputing Center. DT is grateful to the National Science Foundation (CHE 2320256) for support.

