## Supplementary Material for "RNA structural complexity dictates its ion atmosphere"

**Model:** We treated RNAs using the Single Interaction Site (SIS) model where each nucleotide is represented by a single bead, which has been successful in reproducing many aspects of repeat RNAs and mRNAs.[1–4] The divalent cations such as $Mg^{2+}$ and $Ca^{2+}$ are present explicitly in the simulations whereas the monovalent salts are modeled implicitly using the Debye–Huckel approximation. Combining implicit with explicit ion representation facilitates the exploration of RNA configurational space and its ion environment efficiently.[5] We placed the RNA in a cubic box of length from 72 to 90 nm based on the concentration of divalent cations, ensuring at least 200 ions present in the simulation. We applied periodic boundary conditions (PBC) in all 3 dimensions. The effective total energy, $E_{TOT}$, of an RNA-ion system is the sum of bonded ($E_B$), hydrogen bonded ($E_{HB}$), excluded volume ($E_{EV}$), and electrostatic energy ($E_{el}$):

$$E_{TOT} = E_B + E_{HB} + E_{EV}^{RNA} + E_{el}^{P-P} + E_{el}^{X^{2+}-X^{2+}} + E_{EV}^{X^{2+}-X^{2+}} + E_{eff}^{X^{2+}-P}. \tag{S1}$$

The first four terms in Eq. S1 are due to the interactions between RNA beads. The next two terms are for interactions between divalent cations $X^{2+}$ ($Mg^{2+}$ or $Ca^{2+}$), and the last term of the equation is for the effective interactions between $X^{2+}$ and phosphate groups. $E_B$ is the sum of bond stretch ($E_{BS}$) and bond angle ($E_{BA}$) potentials:

$$E_B = E_{BS} + E_{BA} \equiv \frac{1}{2}\sum_{i}^{N_T-1} k_{\text{bond}}(r_i - r_0)^2 + \frac{1}{2}\sum_{j}^{N_T-2} k_{\text{angle}}(\alpha_j - \alpha_0)^2, \tag{S2}$$

where $N_T$ is the number of nucleotides in a given sequence. The parameters $k_{\text{bond}}$ and $k_{\text{angle}}$ are the spring constants in $E_{BS}$ and $E_{BA}$, respectively. $r_i$ and $\alpha_j$ are the $i^{th}$ bond distance and $j^{th}$ bond angle, respectively, with $r_0$ and $\alpha_0$ being their corresponding equilibrium values. The values of all parameters are given in Table S1.

We modeled the hydrogen bond interactions between a pair of nucleotides using:

$$E_{HB} = N_b u_{hb}^0 \exp[-U_{\text{hb,bond}} - U_{\text{hb,angle}} - U_{\text{hb,dihedral}}], \tag{S3}$$

where $N_b = 3$ for a G-C, or $N_b = 2$ for an A-U or G-U base pair (bp), and $u^0_{hb} = -2$ kcal/mol is the strength of a single hydrogen bond, which was calibrated to reproduce the melting temperatures of small CAG repeats.[2]

$U_{\text{hb,bond}}$, $U_{\text{hb,angle}}$ and $U_{\text{hb,dihedral}}$ are calculated as:

$$U_{\text{hb,bond}} = k_r(r_{ij} - r_{hb,0})^2, \tag{S4}$$

$$U_{\text{hb,angle}} = k_\theta(\theta_{i,j,j-1} - \theta_1)^2 + k_\theta(\theta_{i-1,i,j} - \theta_1)^2 + k_\theta(\theta_{i,j,j+1} - \theta_2)^2 + k_\theta(\theta_{i+1,i,j} - \theta_2)^2, \tag{S5}$$

$$U_{\text{hb,dihedral}} = k_\phi[1 + cos(\phi_{j-1,j,i,i-1} + \phi_1)] + k_\phi[1 + cos(\phi_{j+1,j,i,i+1} + \phi_2)]. \tag{S6}$$

In Eq. S4, $r_{ij}$ is the distance between bead $i$ and its complementary bead $j$, $r_{hb,0}$ is the equilibrium distance in an ideal A-form helix. In Eq. S5, $\theta_{i,j,k}$ is the angle between beads $i$, $j$ and $k$. The contribution of the angular part to the total hydrogen bond potential is determined by $k_\theta$. In Eq. S6, the dihedral angle, $\phi_{i,j,k,l}$, is the angle between the two planes defined by the four beads $i$, $j$, $k$ and $l$.

To prevent unphysical overlaps between two non-bonded RNA beads, we applied the Weeks–Chandler–Andersen repulsive potential given as,

$$E^{RNA}_{EV} = \epsilon_{ev}\left[\left(\frac{\sigma}{r_{ij}}\right)^{12} - 2\left(\frac{\sigma}{r_{ij}}\right)^{6} + 1\right] \text{ if } r_{ij} \le \sigma, \tag{S7}$$

$$E^{RNA}_{EV} = 0 \text{ otherwise,} \tag{S8}$$

where $\epsilon_{ev} = 2.0$ kcal/mol is the strength of the repulsive interactions, and $\sigma = 10$ Å is the diameter of an RNA bead. We applied the same potential between two ions, $E^{X^{2+}-X^{2+}}_{EV}$, (X = Ca or Mg). The values of $\epsilon_{ev}$ for ions and their diameters $\sigma^{X^{2+}}$ are taken from Nguyen et al,[5] (listed in Table S1).

**Effective $X^{2+}$-P interactions, $E^{X^{2+}-P}_{eff}$:** Following our previous work,[5] we used a combination of short ranged interactions derived from the Reference Interaction Site Model (RISM) theory and long ranged electrostatic interactions from the Debye–Huckel approximation to account for the effective interactions between a divalent cation, $X^{2+}$, and a phosphate group:

$$E_{eff}^{X^{2+}-P} = W(r) + [U_{DH}(r) - W(r)] \exp\left(-\frac{a^2}{r^2}\right), \tag{S9}$$

where $W(r)$ is the potential of mean force (PMF) derived from RISM theory, and $U_{DH}(r)$ is the Debye–Huckel interaction accounting for the screening effects of monovalent salts. We set $a = 5$ Å to preserve the minimum value for $X^{2+}$-P interaction energy. It is important to note that $E_{eff}^{X^{2+}-P}$ is dominated by $W(r)$ if $r$ is small. At large separation, $U_{DH}(r)$ is dominant. The tabulated values of $W(r)$ were taken from Nguyen et al.[5]

**Langevin Dynamics Simulations:** To efficiently explore the conformational ensembles of RNAs, we performed low-friction Langevin dynamics simulations at temperature $T = 27^{\circ}$ C using "LangevinIntegrator" in OpenMM.[6] The Langevin dynamics equation of motion is given by:

$$m_i \frac{\mathrm{d}\mathbf{v_i}}{\mathrm{d}t} = \mathbf{f_i} - \gamma m_i \mathbf{v_i} + \mathbf{R_i}, \tag{S10}$$

where $m_i$ and $\mathbf{v_i}$ are the mass and velocity of the particle $i$, respectively, $\mathbf{f_i}$ is the deterministic force acting on particle $i$, and $\mathbf{R_i}$ is an uncorrelated random force on particle $i$ whose components are chosen from a normal distribution with zero mean and unit variance. We set the friction coefficient $\gamma = 1$ ps$^{-1}$. We integrated the equation of motion using a time step $\delta t = 2$ fs.

**Brownian Dynamics Simulations:** We also performed Brownian dynamics simulations at $T = 27^{\circ}$ C with a friction coefficient equal to that of water using "BrownianIntegrator" in OpenMM to study the dynamical properties of $Mg^{2+}$ ions. The Brownian dynamics equation of motion is given by:

$$\frac{\mathrm{d}\mathbf{r_i}}{\mathrm{d}t} = \frac{1}{\gamma m_i}\mathbf{f_i} + \mathbf{R_i}, \tag{S11}$$

where $\mathbf{r_i}$ and $\mathbf{f_i}$ are the position and force acting on the particle $i$, respectively. $\gamma$ ($\approx 91$ ps$^{-1}$) is the friction coefficient of water. $\mathbf{R_i}$ is the uncorrelated random force on the particle $i$ whose components are chosen from a normal distribution with zero mean and variance equal to $2k_{\mathrm{B}}T/\gamma m_i$. We used $\delta t = 5$ fs for those simulations.

**Table S1: List of parameters used in the single bead SOP energy function**

| | |
|---|---|
| $k_{bond}$ | 15 kcal/mol. Å$^{-2}$ |
| $r_0$ | 5.9 Å |
| $k_{angle}$ | 3.0 kcal/mol.rad$^{-2}$ |
| $\alpha_0$ | 2.618 rad |
| $\epsilon_{EV}$ | 2.0 kcal/mol |
| $\epsilon_{EV}^{Mg^{2+}}$ | 0.895 kcal/mol |
| $\epsilon_{EV}^{Ca^{2+}}$ | 1.000 kcal/mol |
| $\sigma$ | 10 Å |
| $\sigma^{Mg^{2+}}$ | 4 Å |
| $\sigma^{Ca^{2+}}$ | 5.6 Å |
| $r_{hb,0}$ | 13.8 Å |
| $k_r$ | 3.0 Å$^{-2}$ |
| $k_\theta$ | 1.5 rad $^{-2}$ |
| $k_\phi$ | 0.5 |
| $\theta_1$ | 1.8326 rad |
| $\theta_2$ | 0.9425 rad |
| $\phi_1$ | 1.8326 rad |
| $\phi_2$ | 1.345 rad |
| $u_{hb,0}$ | -2.0 kcal/mol |

**RNA structural heterogeneity in monovalent salts:** To assess the RNA structural ensemble, we performed simulations of RNAs in implicit monovalent salt, which is modeled using the Debye–Huckel approximation[7] (concentration ranges between 20 mM and 150 mM) (Fig. S1). The average radius of gyration, $R_g^{sim}$, decreases as the concentration increases from 20 mM to 150 mM, which is in good agreement with experimental SAXS measurements.[8]

The calculated average distance for a pair of residues $i$ and $j$, $\langle R_{ij} \rangle$, as a function of the separation length $|i-j|$, indicates compaction at high salt concentrations is caused by long-ranged interactions ($|i-j| \geq 8$) (Fig. S1B). However, at those conditions, $rU_{30}$ is expanded due to strong electrostatic repulsions between phosphate groups. The computed contact probability $\langle Q_{ij} \rangle$, with respect to a Gaussian polymer chain, is featureless and bears the signature of a heterogeneous and disordered conformational ensemble (Fig. S1C - S1F).

Unlike $rU_{30}$, $\langle Q_{ij} \rangle$ of the CAG repeat exhibits a few strong contacts and bears the signature of typical stem-loop like configurations. However, the presence of multiple contact points for a given nucleotide $i$ indicates a heterogeneous ensemble for the CAG repeat. Moreover, we observed a substantial increase of $\langle Q_{ij} \rangle$ with an increase in salt concentrations (Fig. S3) due to the enhanced electrostatic screening effect. To quantify the stem-loop configurations, we calculated the joint probability distribution of $Q_{HP}$ and $N_{bp}$, $P(Q_{HP}, N_{bp})$. When the ion concentration increases from 20 mM to 150 mM, the maximum population of hairpins shifted from $Q_{HP} = 5$ and $N_{bp} = 2$ to $Q_{HP} = 1$ and $N_{bp} = 6$ (Fig. S5).

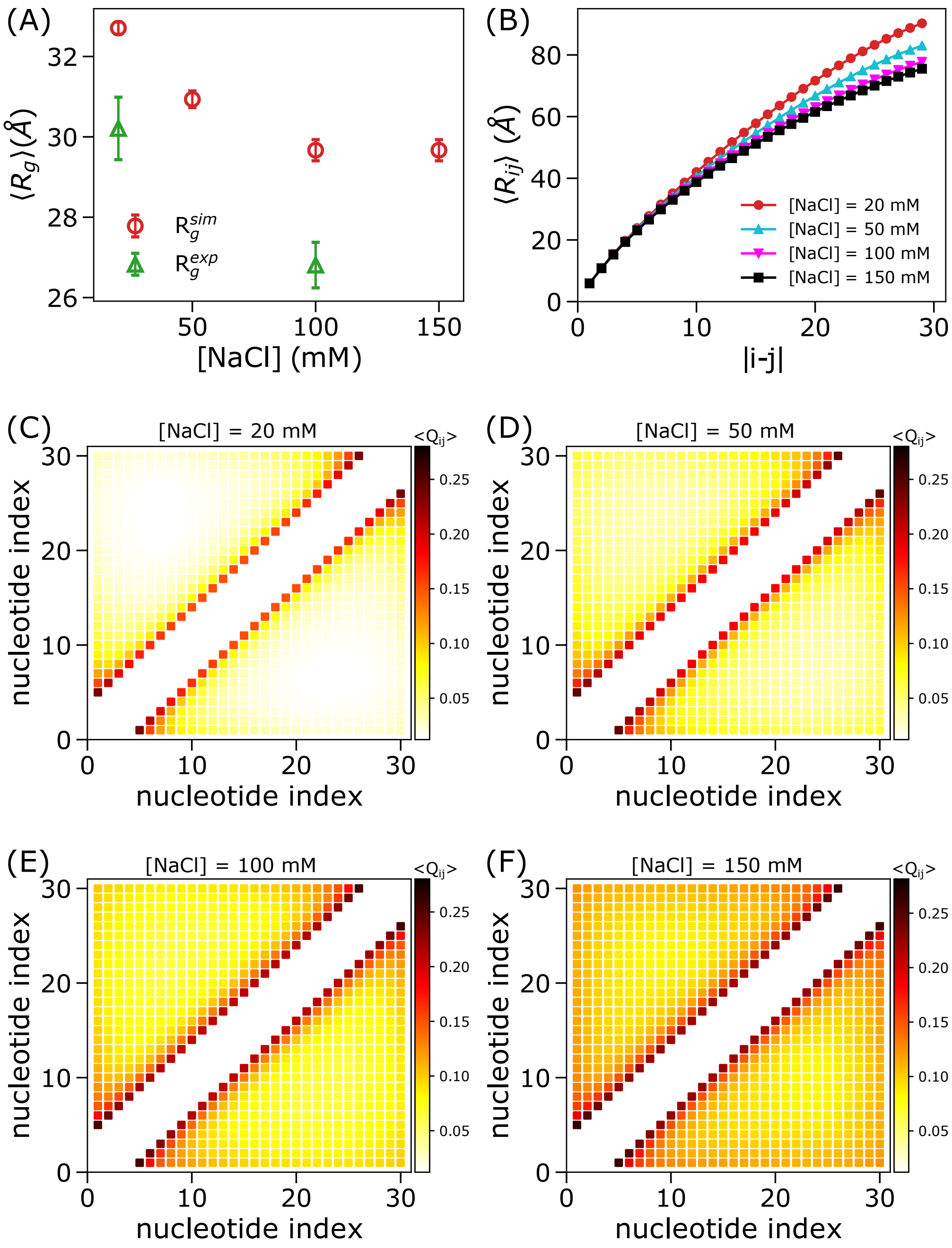


Figure S1: Structural ensemble of rU$_{30}$ in monovalent salt: (A) The simulated and experimentally[8] determined radius of gyration, $\langle R_g \rangle$, denoted as $R_g^{sim}$ (red) and $R_g^{exp}$ (green), respectively, are shown as a function of monovalent salt concentration. The large error bars associated with the data points reflect significant conformational heterogeneity in the ensemble. (B) Average distance, $\langle R_{ij} \rangle$, as a function of segment length, $|i-j|$, at 20 mM (red), 50 mM (cyan), 100 mM (magenta) and 150 mM (black) salt concentration indicates compaction is driven by interactions between nucleotides with segment length $> 10$. (C–F) Simulated contact probability, $\langle Q_{ij} \rangle$, between nucleotides $i$ and $j$ at different salt concentrations: (C) 20 mM, (D) 50 mM, (E) 100 mM, and (F) 150 mM. The increase in $\langle Q_{ij} \rangle$ with rising salt concentration reflects reduced nucleotide-nucleotide repulsion due to stronger screening effects at higher salt concentrations.

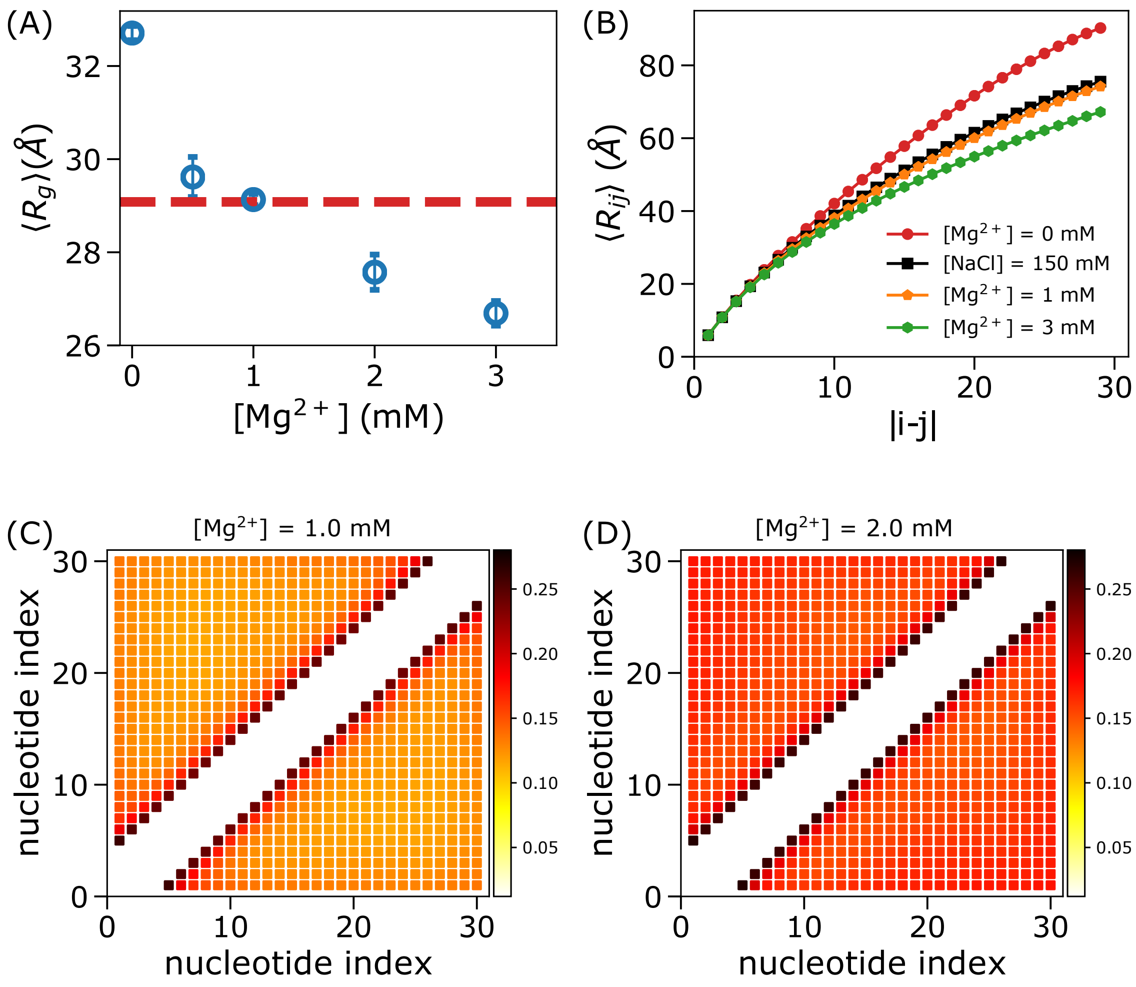


Figure S2: Effect of $Mg^{2+}$ on $rU_{30}$ structure: (A) RNA compaction at increasing $Mg^{2+}$ concentrations. (B) Average distance, $\langle R_{ij} \rangle$, as a function of sequence separation length, $|i-j|$, between two nucleotides $i$ and $j$ at 20 mM NaCl + 0.0 mM (red), or 1.0 mM (orange), or 3.0 mM (green) $Mg^{2+}$ indicates compaction is driven by long-range interactions. Also the data at 150 mM NaCl with no $Mg^{2+}$ (black) is shown for comparison. (C) $\langle Q_{ij} \rangle$ at 20 mM NaCl + 1.0 mM $Mg^{2+}$. (D) Same as C, but at 2.0 mM $Mg^{2+}$.

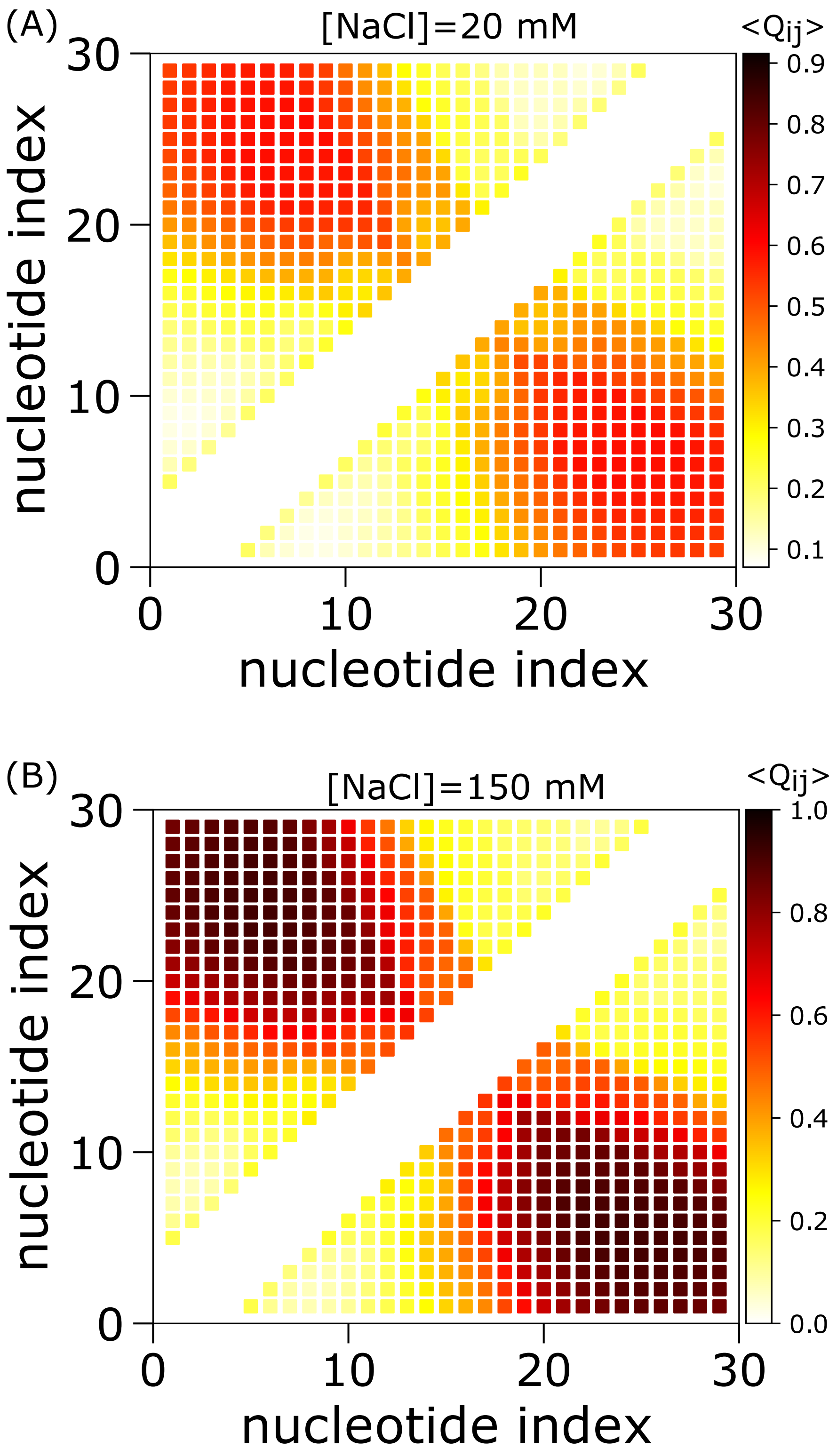


Figure S3: Average contact probability, $\langle Q_{ij} \rangle$, between two nucleotides, $i$ and $j$ for $A(CAG)_9A$ in 20 mM and 150 mM monovalent salt.

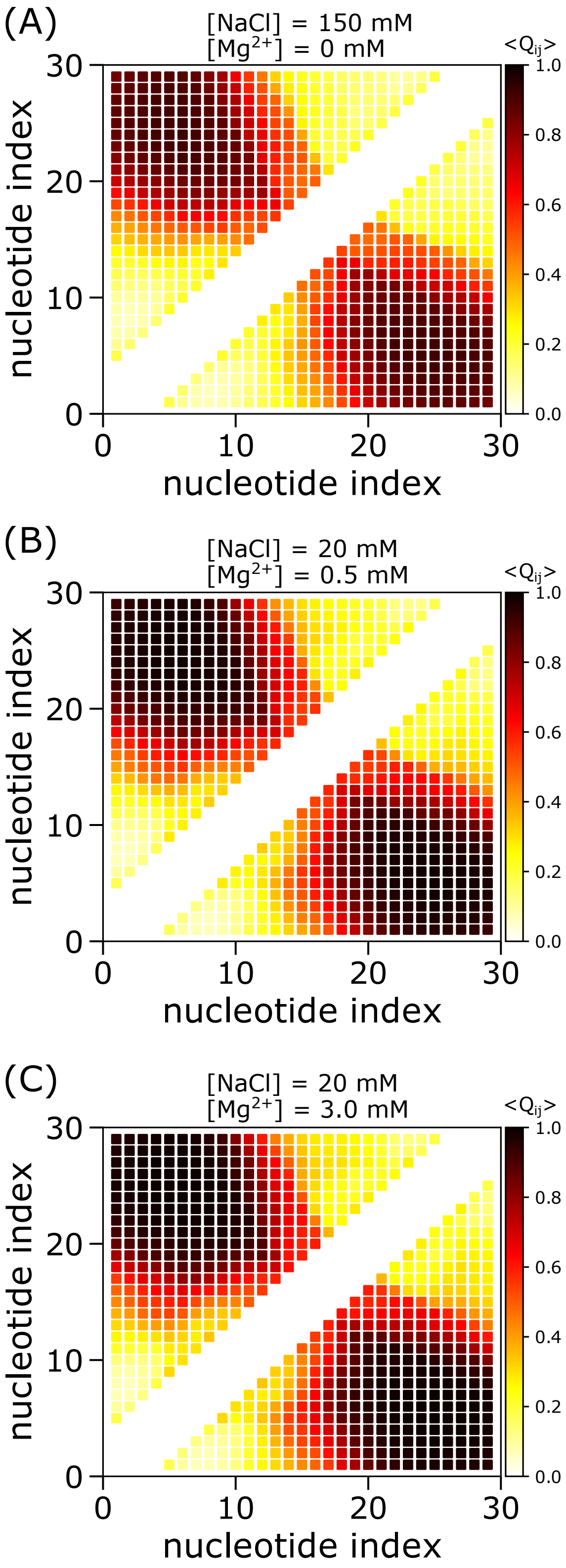


Figure S4: $Q_{ij}$ for A(CAG)$_9$A at varying ion conditions.

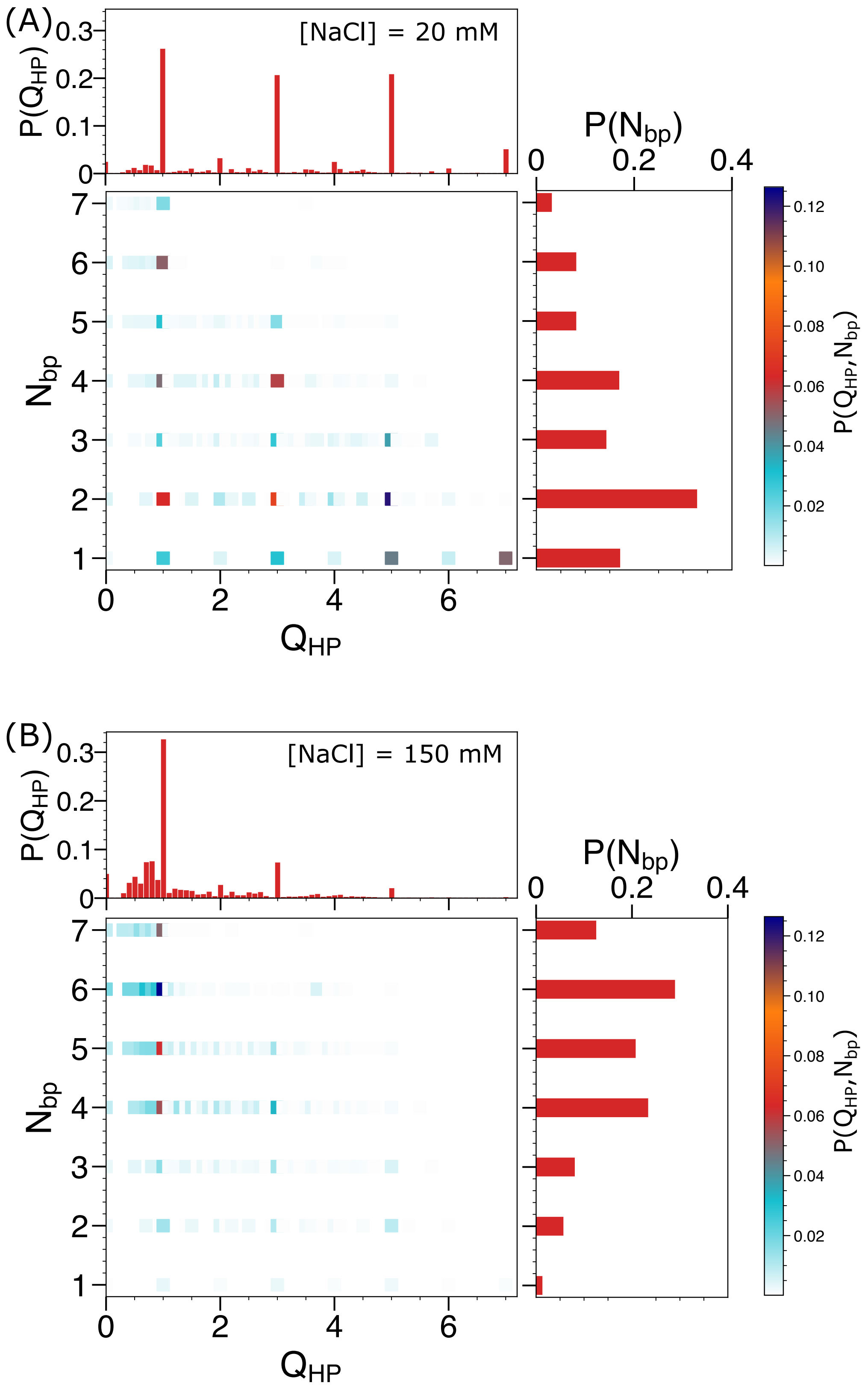


Figure S5: Joint probability distribution of $Q_{HP}$ and $N_{bp}$, $P(Q_{HP}, N_{bp})$, for A(CAG)$_9$A in monovalent salt: (A) At 20 mM. (B) At 150 mM. The distributions of $Q_{HP}$ ($P(Q_{HP})$) and $N_{bp}$ ($P(N_{bp})$) are in top and right panels, respectively.

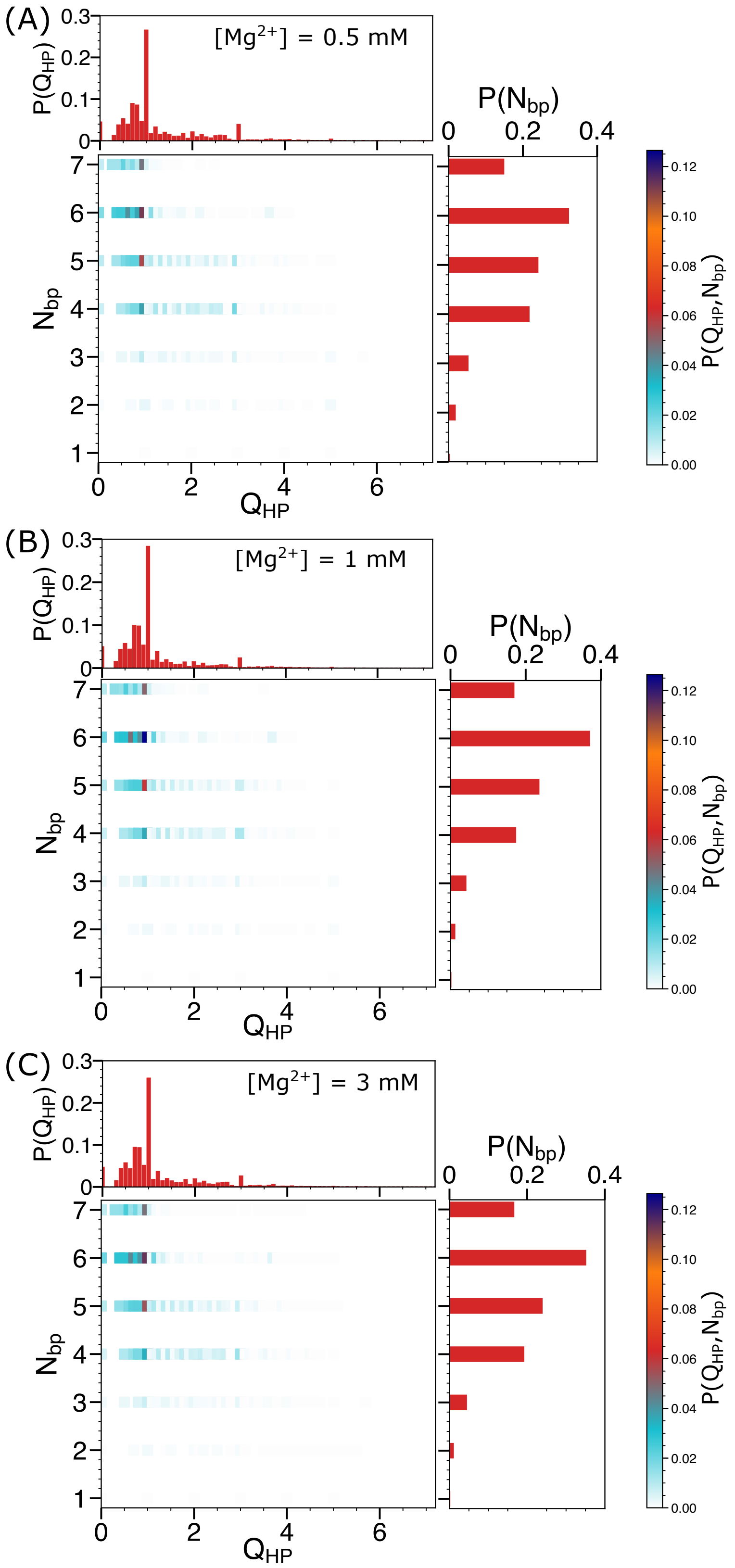


Figure S6: Joint probability distribution of $Q_{HP}$ and $N_{bp}$ for A(CAG)$_9$A at increasing [Mg$^{2+}$].

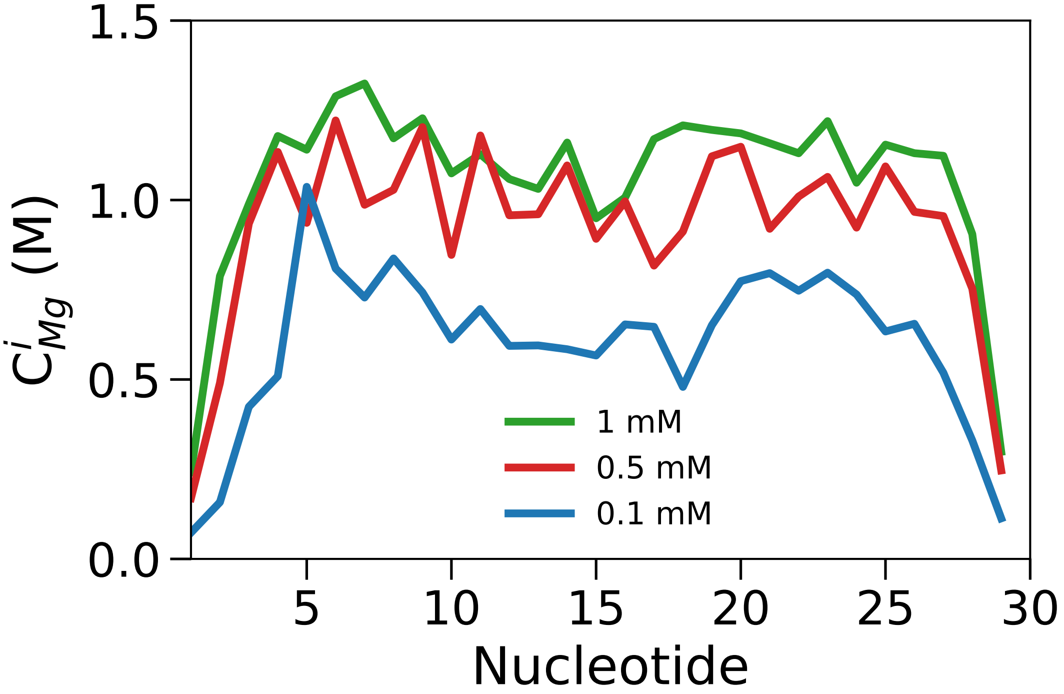


Figure S7: Local ion concentration of $Mg^{2+}$ around individual nucleotides for $A(CAG)_9A$ at 1 mM (green), 0.5 mM (red) and 0.1 mM (blue) $Mg^{2+}$ concentrations.

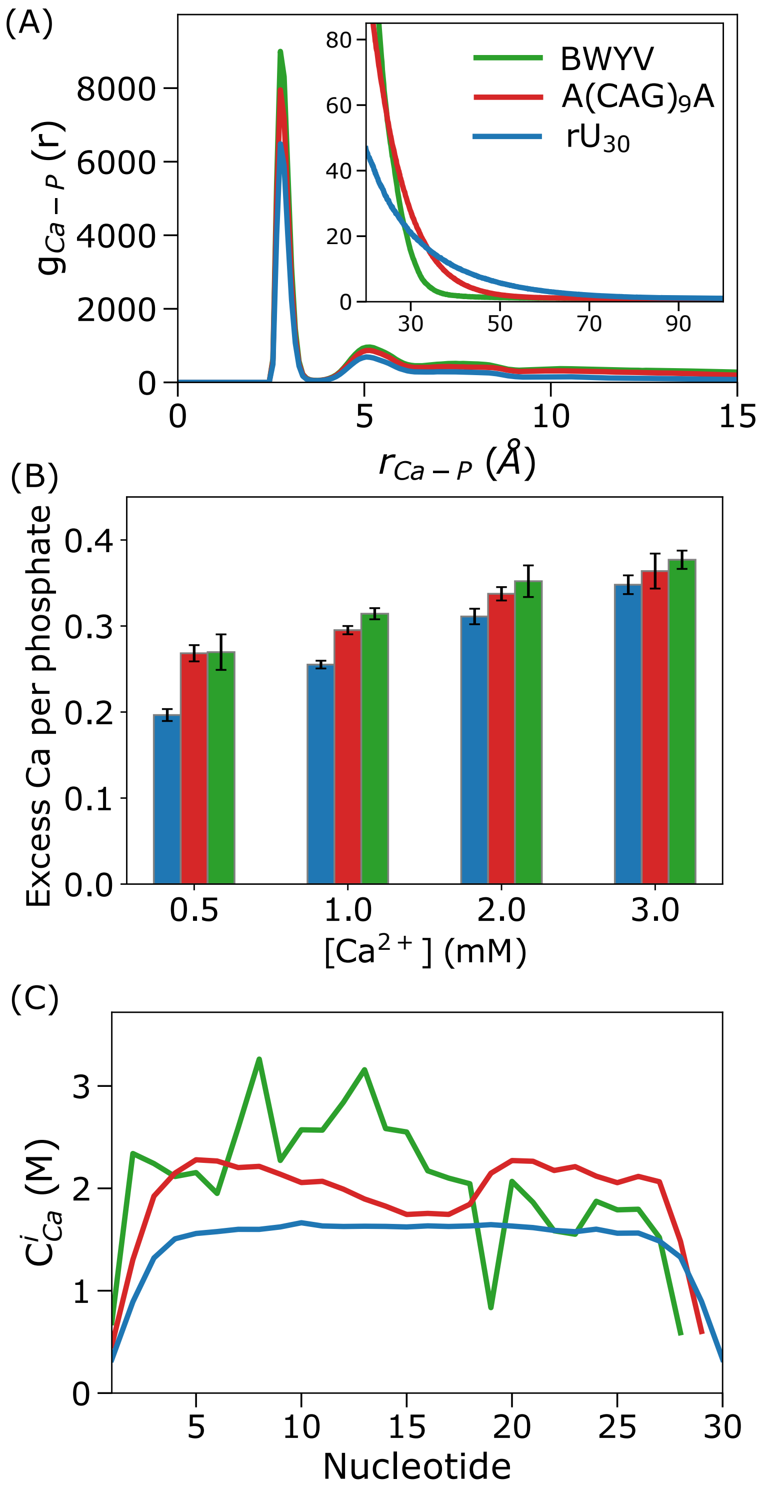


Figure S8: $Ca^{2+}$ distribution around RNAs: (A) Radial distribution function $g_{\mathrm{Ca-P}}(r)$ between $Ca^{2+}$ and phosphate groups for $rU_{30}$, $A(CAG)_9A$ and BWYV. The inset indicates $g_{\mathrm{Ca-P}}(r)$ decays rapidly for well-structured RNAs. (B) Excess $Ca^{2+}$ per phosphates for $rU_{30}$, $A(CAG)_9A$ and BWYV are in blue, green and red, respectively. (C) Local $[Ca^{2+}]$ indicates the ion concentration is correlated with the phosphate density. All calculations were performed at 20 mM NaCl + 0.5 mM $Ca^{2+}$.

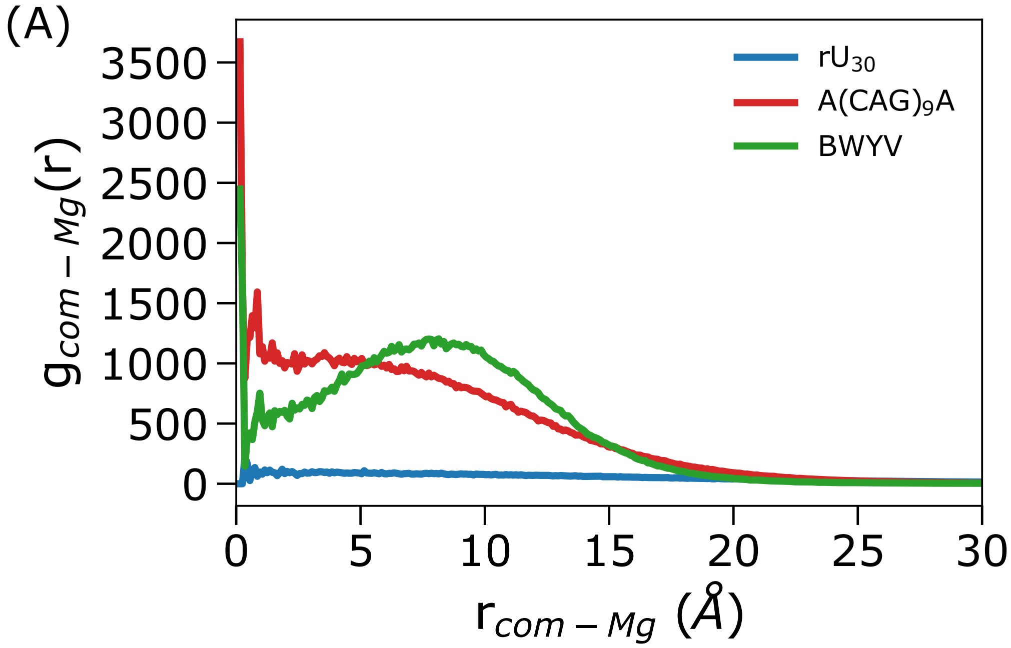


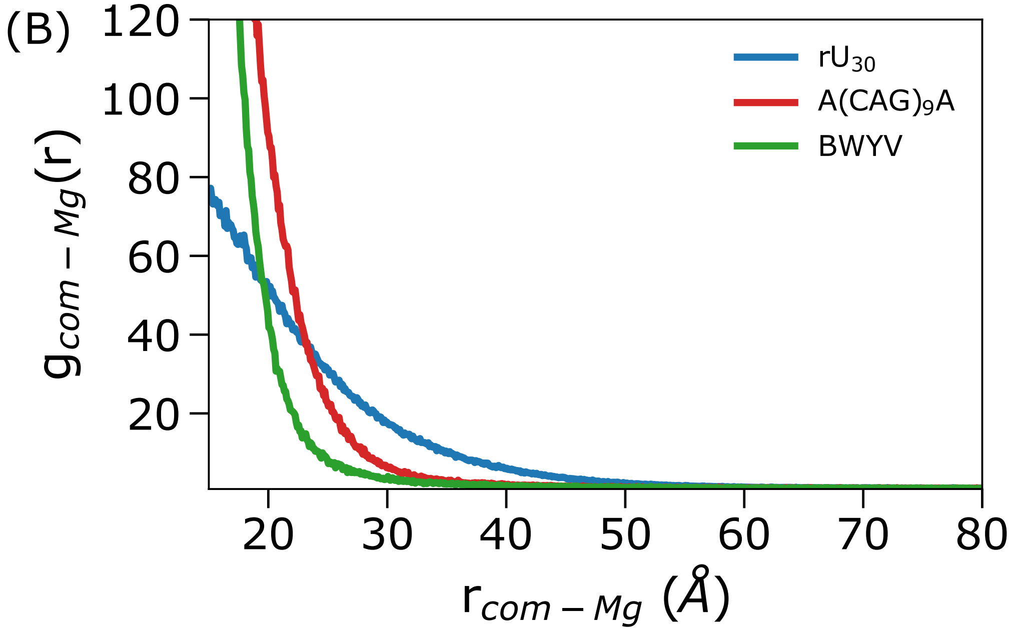


Figure S9: Radial distribution function of $Mg^{2+}$ ions with respect to the center of mass of the RNAs.

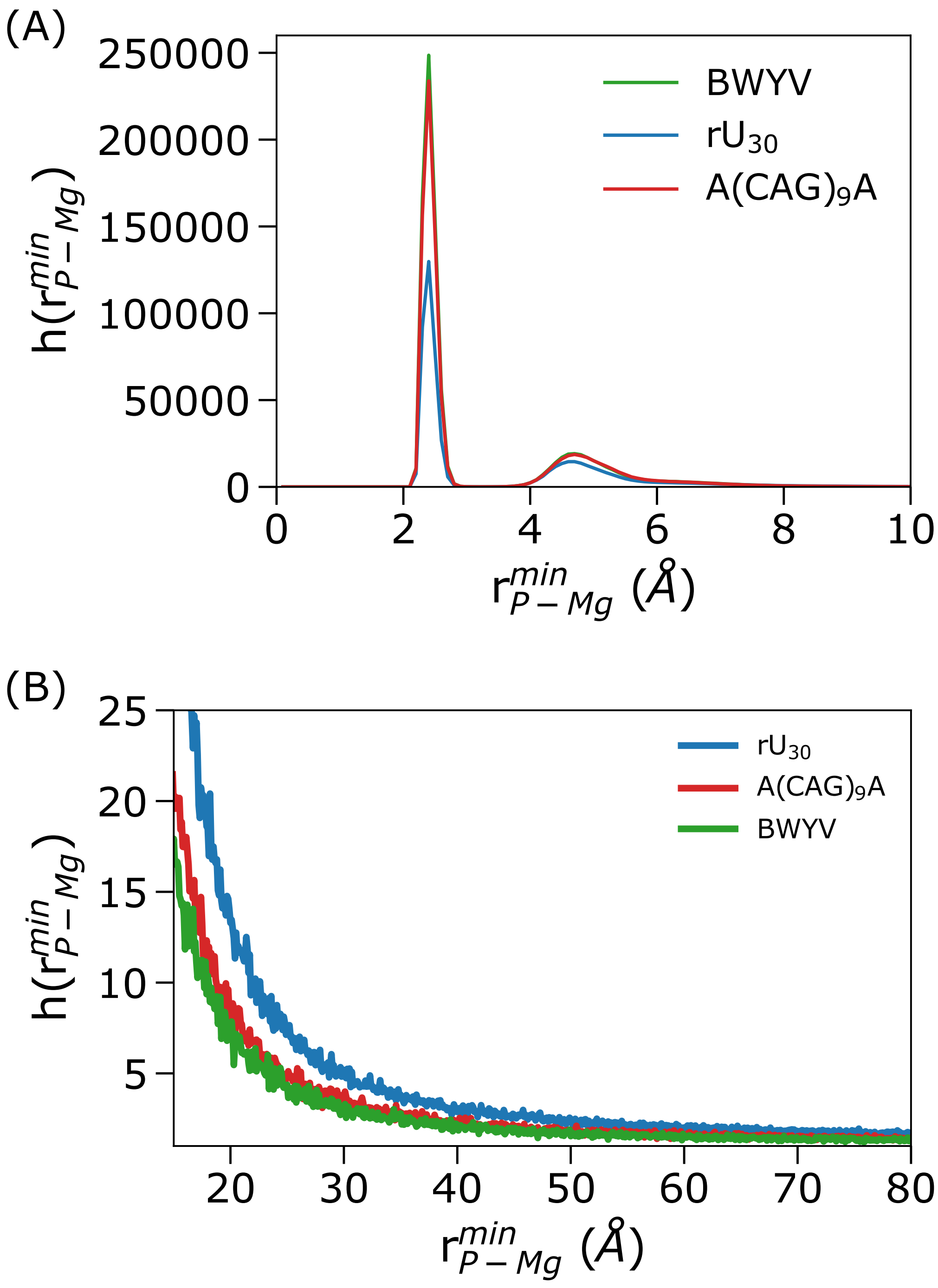


Figure S10: Histogram of $Mg^{2+}$ ions to the closest phosphate group of RNAs.

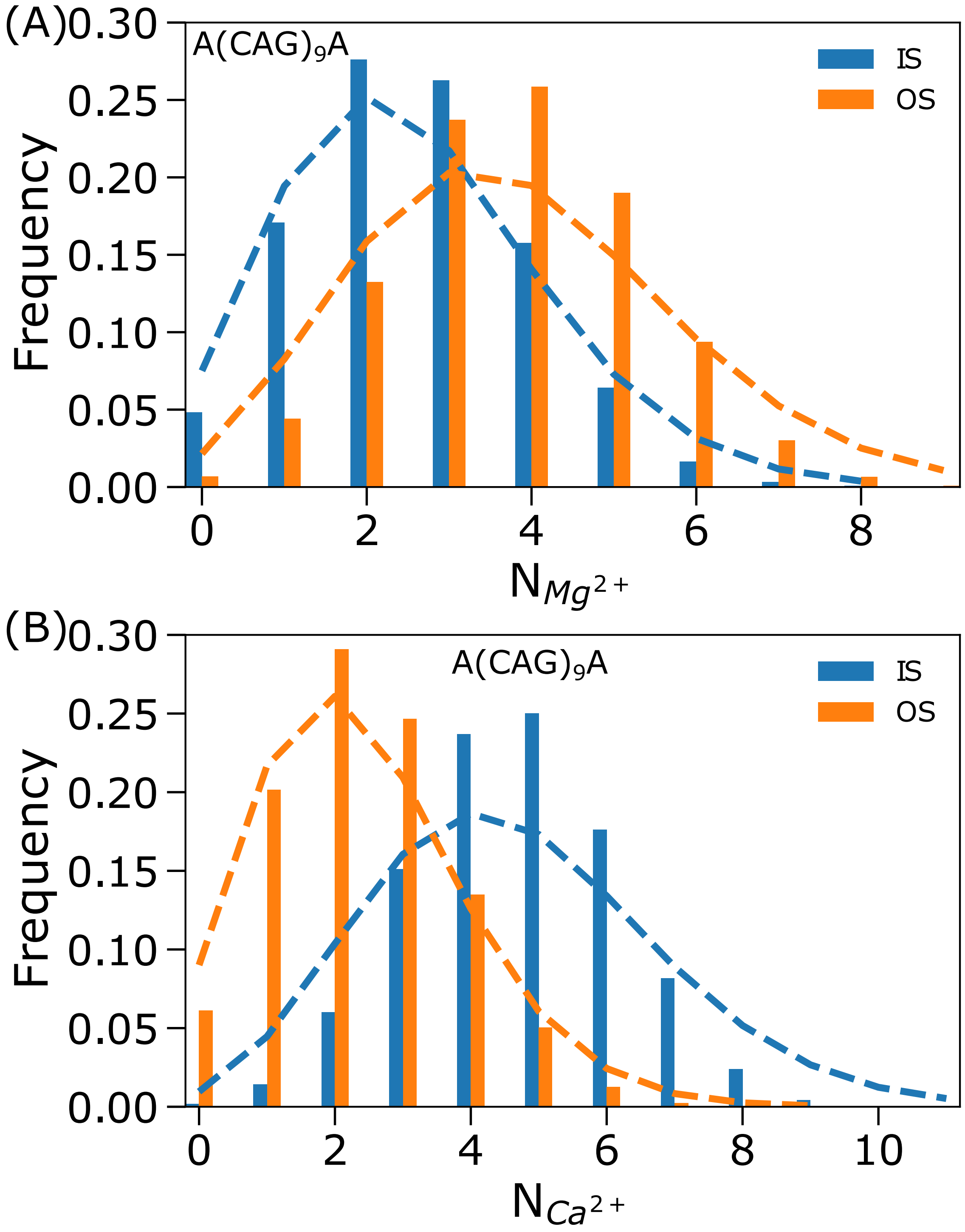


Figure S11: Distribution of inner and outer sphere ions around the CAG repeat: (A) Probability distributions of inner and outer sphere $Mg^{2+}$ ions around $A(CAG)_9A$ are in blue and orange, respectively. The broken lines are obtained from Poisson distributions using the mean values of inner and outer sphere ions, respectively. (B) Same as (A) except for $Ca^{2+}$ ions.

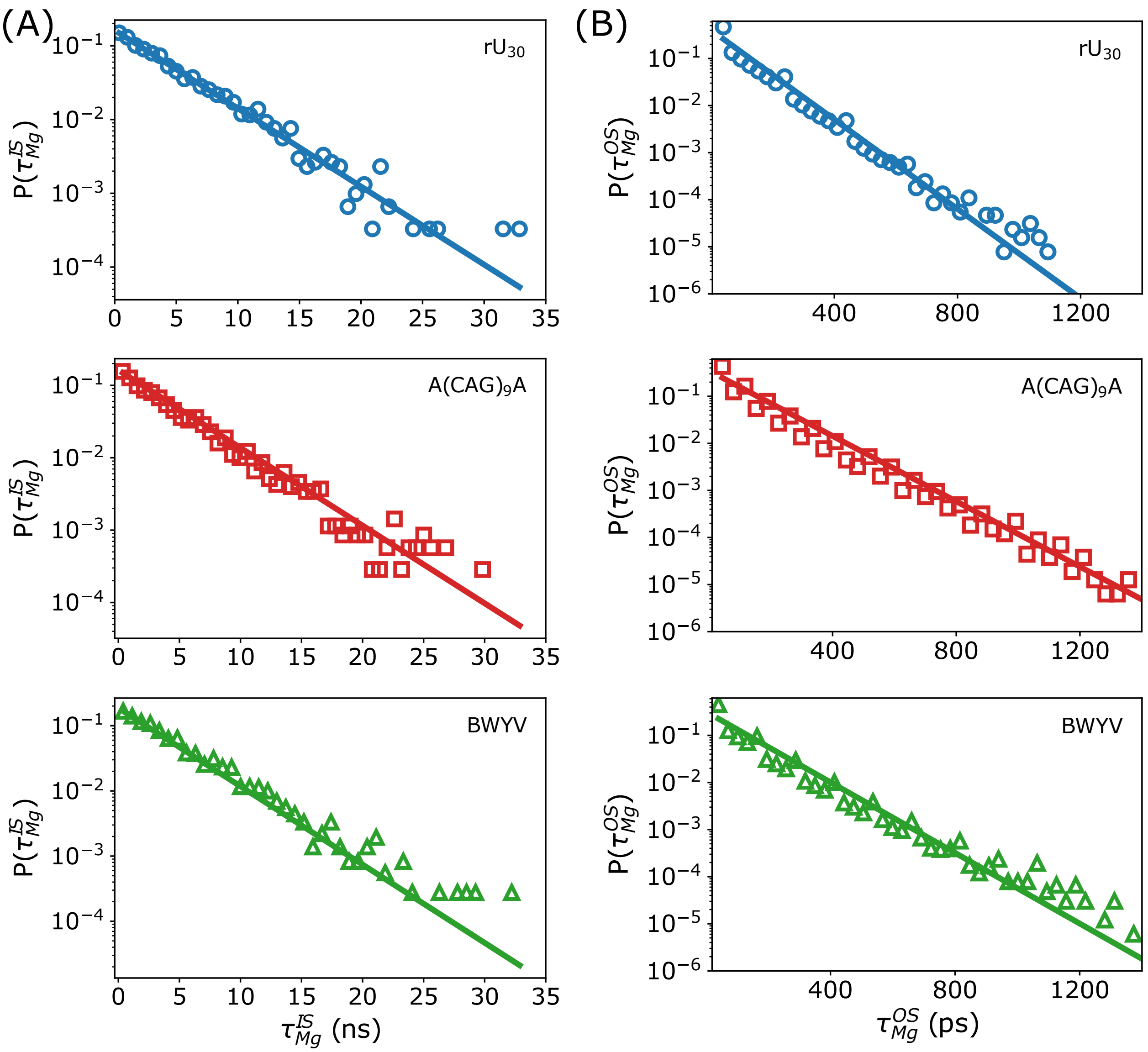


Figure S12: Dwell times of IS (A) and OS (B) $Mg^{2+}$ ions follow a single exponential decay for $rU_{30}$ (top), $A(CAG)_9A$ (middle) and BWYV (bottom).

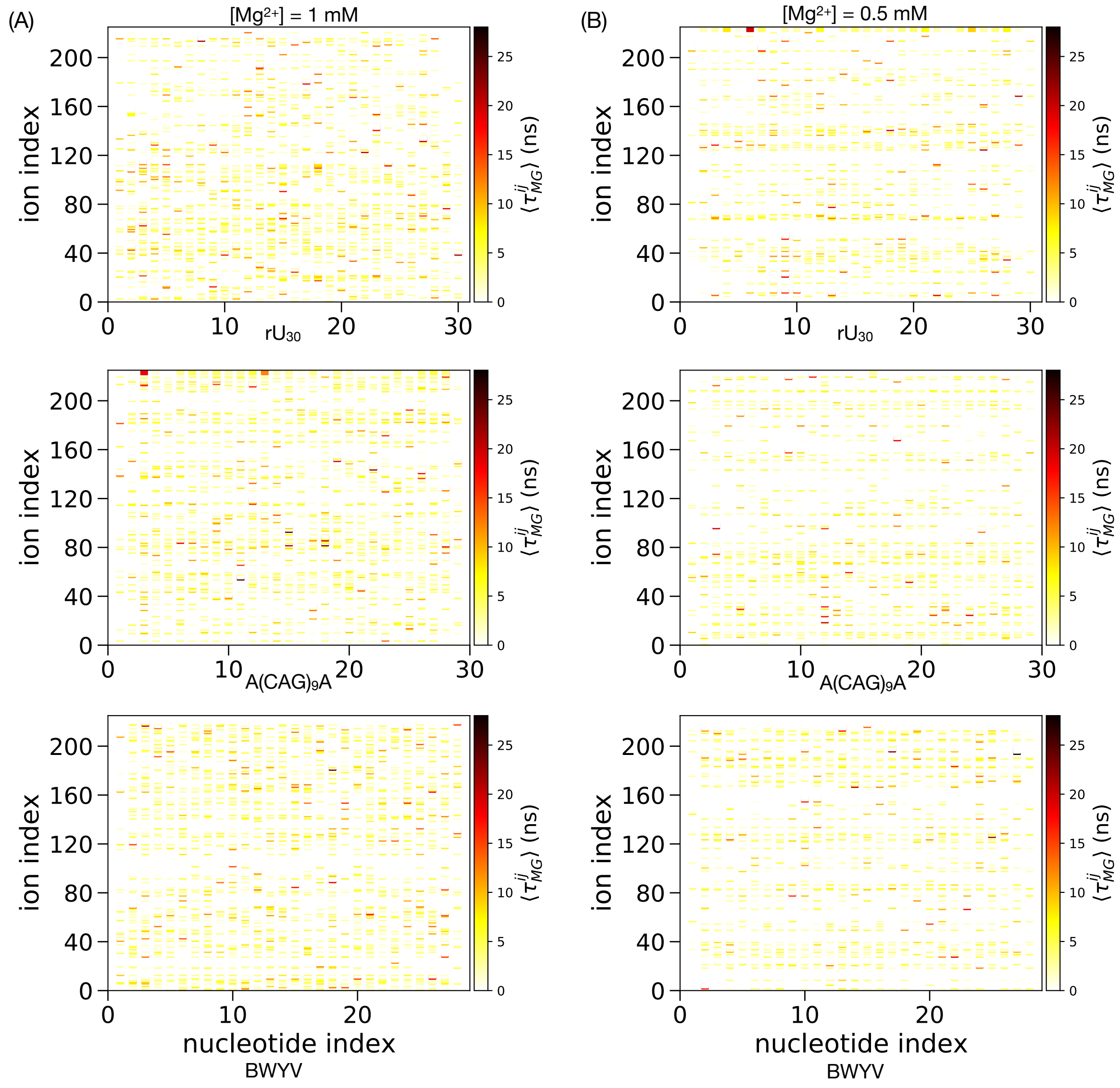


Figure S13: Average dwell time of ion $j$ around nucleotide $i$: (A) $\langle \tau_{Mg}^{ij} \rangle$ at 1.0 mM $Mg^{2+}$ for $rU_{30}$ (top), $A(CAG)_9A$ (middle) and BWYV (bottom). (B) Same as A, but at 0.5 mM $Mg^{2+}$.

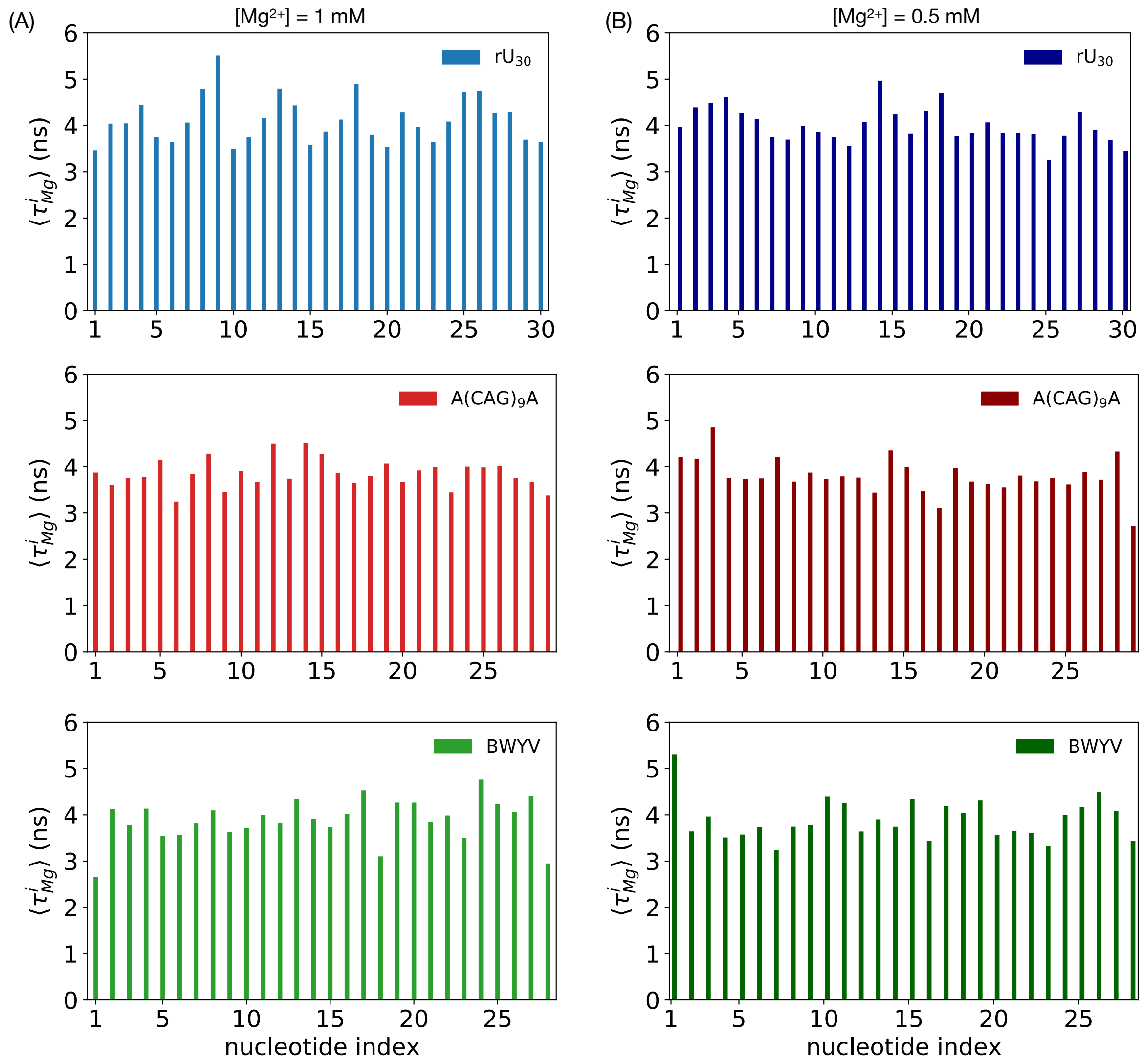


Figure S14: Average dwell time of $Mg^{2+}$ around nucleotide $i$: (A) $\langle \tau^i_{Mg} \rangle$ at 1.0 mM $Mg^{2+}$ for $rU_{30}$ (top), $A(CAG)_9A$ (middle) and BWYV (bottom). (B) Same as A, but at 0.5 mM $Mg^{2+}$.
